# Kif2C safeguards radial glial integrity to prevent cortical malformation

**DOI:** 10.64898/2026.05.22.726143

**Authors:** Sharmin Naher, Takako Kikkawa, Kenji Iemura, Haw-Yuan Cheng, Shinsuke Niwa, Amjad Khan, Magdalena Krygier, Michael Zech, Meng-Han Tsai, Maria Mazurkiewicz-Bełdzińska, Jin-Wu Tsai, Laurent Nguyen, Kozo Tanaka, Noriko Osumi

## Abstract

Radial glial cells (RGCs) generate cortical neurons and guide radial neuronal migration, yet how microtubule (MT) regulators coordinate progenitor maintenance, mitotic fidelity, and cortical architecture remains unclear. Here, we identify the MT depolymerase Kif2C/MCAK as an essential regulator of RGC integrity with relevance to human neurodevelopmental disorders. Kif2C is enriched in developing cortical RGCs and localizes to radial fibers, basal endfeet, and mitotic structures. Acute Kif2C depletion in embryonic mouse cortices disrupts RG fiber organization, impairs neuronal migration, reduces the RGC pool, and induces mitotic defects, chromosome segregation errors, DNA damage, and cell-cycle arrest. *Kif2C*-deficient cortices further exhibit focal pial basement membrane disruption and neuronal overmigration, resulting in a cobblestone-like cortical malformation. We identify two individuals with neurodevelopmental disorders carrying rare deleterious *KIF2C* variants and show that a patient-derived truncating variant fails to rescue *Kif2C*-deficient cortical phenotypes, implicating KIF2C dysfunction in human neurodevelopmental disorders.

**Graphical abstract:** 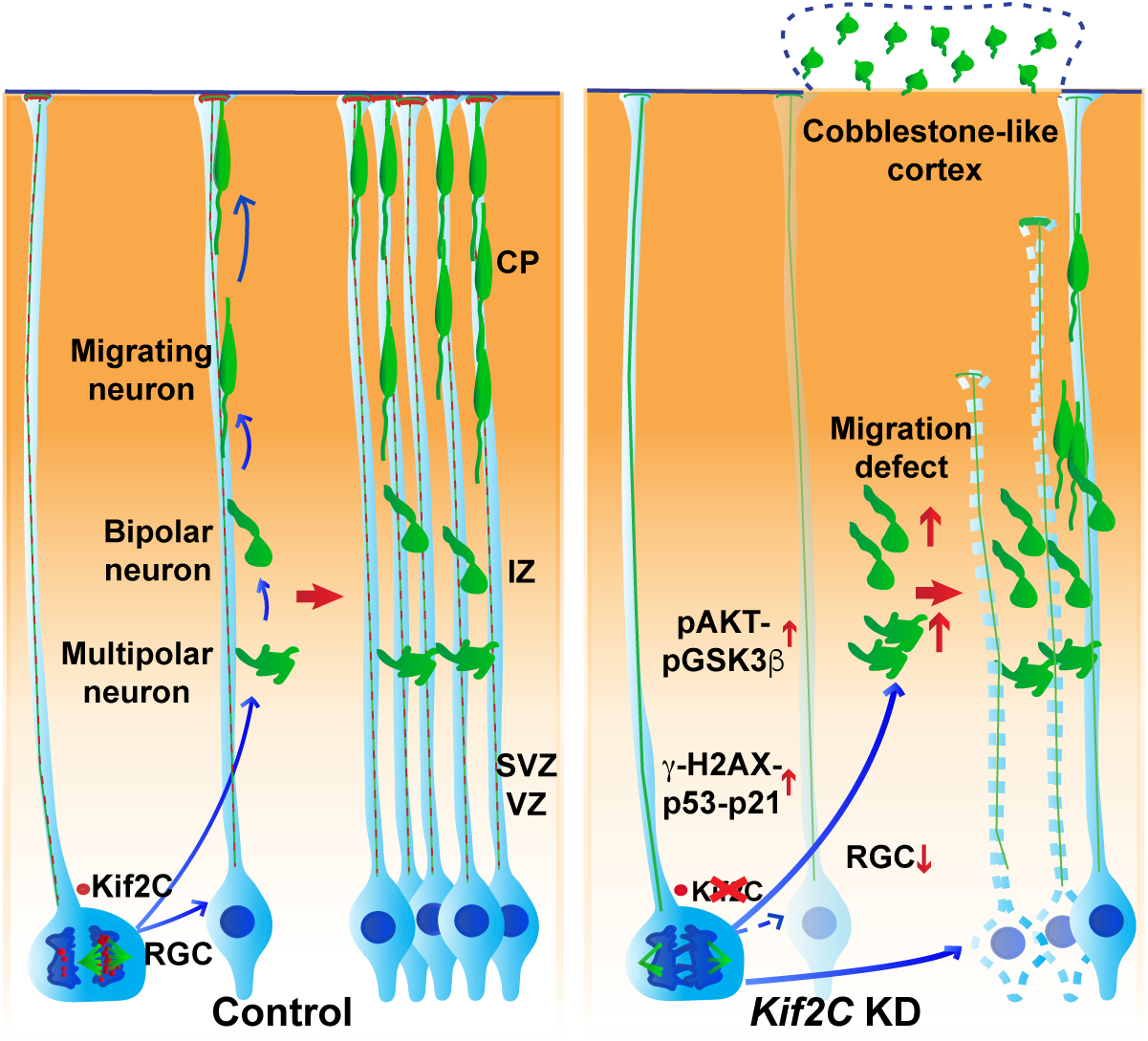

## Introduction

During cortical development, radial glial cells (RGCs) play two essential roles: they serve as the primary neural progenitors of the cerebral cortex and provide the radial scaffold that guides the migration of newborn excitatory neurons. RGC cell bodies reside in the ventricular zone (VZ), whereas their long basal fibers extend toward the pial surface and terminate as specialized endfeet that contact the pial basement membrane [1]. This highly polarized architecture is crucial not only for neuronal production and radial migration but also for maintaining the structural boundary of the developing cortex [2,3,4]. Disruptions of RGC proliferation, morphology, or endfoot attachment can therefore perturb multiple aspects of cortical development simultaneously, resulting in abnormal neuronal positioning, laminating defects, and malformations of cortical development [5,6]. In particular, cobblestone malformations, which are characterized by breaches of the pial basement membrane and neuronal overmigration beyond the cortical surface [7], highlight the importance of mechanisms that preserve radial scaffold integrity and the pial boundary.

A central cellular system underlying these RGC functions is the microtubule (MT) cytoskeleton. MTs and MT-associated proteins are required for mitotic spindle assembly, chromosome segregation, and interkinetic nuclear migration of cortical progenitors, and they also support the elongated morphology of RG fibers [8,9]. Among the major regulators of MT dynamics are kinesin-13 family proteins, including Kif2A, Kif2B, and Kif2C, which act as ATP-dependent MT depolymerases rather than cargo-transporting motors [10,11]. MT regulation is clearly essential for both progenitor division and RG architecture; how an MT depolymerase coordinates these dual demands during cortical development remains poorly understood.

Kif2C, also known as mitotic centromere-associated kinesin (MCAK), is a prototypical kinesin-13 family member best known for its role in mitosis. In proliferating cells, Kif2C regulates kinetochore-MT attachments and spindle dynamics to ensure accurate chromosome congression and segregation [12,13]. Beyond mitosis, Kif2C has also been implicated in MT-dependent remodeling during cell migration [14,15], as well as in neuronal contexts such as synaptic structure and plasticity [16]. Notably, while this work was in progress, a recent preprint reported that *Kif2C* knockout mice exhibit cortical disorganization and neuronal defects [17], further supporting the importance of Kif2C in corticogenesis. However, the cellular basis of these developmental abnormalities remains unclear, and whether Kif2C acts in RGCs to coordinate cytoskeletal organization and mitotic fidelity has not been investigated. This question is particularly important because RGCs must simultaneously maintain a highly elongated polarized scaffold and execute error-free proliferative divisions, two processes that are both critically dependent on MT control.

Here, we identify Kif2C as a key regulator of RG integrity during corticogenesis. We show that Kif2C is enriched in mouse cortical RGCs, localizes to radial fibers, basal endfeet, and mitotic structures, and confirm KIF2C protein expression in RG-like cells of the developing human cortex. Acute *Kif2C* depletion in the developing mouse cortex disrupts RG fiber organization, impairs neuronal migration, reduces RGC pool, and causes mitotic abnormalities associated with chromosome segregation defects and activation of a DNA damage-linked cell-cycle arrest program. Strikingly, Kif2C-deficient cortices exhibit focal breaches of the pial basement membrane and neuronal overmigration, resulting in a cobblestone-like malformation phenotype. Finally, we identify two damaging KIF2C variants in unrelated individuals with overlapping neurodevelopmental disorder phenotypes, including microcephaly, intellectual disability, global developmental delay, and motor impairment. Functional rescue experiments further show that, unlike wild-type human *KIF2C,* the patient-derived p.Trp230* variant fails to restore RG fiber integrity after mouse *Kif2C* knockdown. Together, these findings establish Kif2C as a factor that links RG cytoskeletal organization and mitotic integrity to cortical tissue architecture and support impaired KIF2C function as a contributor to human neurodevelopmental pathology.

## Results

### Kif2C is expressed in radial glial cells within the developing neocortex

To investigate the potential role of Kif2C in neocortical development, we first examined its expression pattern in the embryonic mouse cortex. Reanalysis of a published single-cell RNA sequencing (scRNA-seq) dataset from embryonic day (E) 14.5 cortex [18] revealed that Kif2C expression is enriched in radial glial cells (RGCs), particularly in a proliferative RGC cluster marked by Top2a expression (Figures 1A and S1). Consistently, analysis of a spatial transcriptomic dataset from the E15.5 mouse brain [19] showed that Kif2C expression is largely confined to the ventricular zone (VZ), where RGCs reside (Figure 1B).

**Figure 1.**
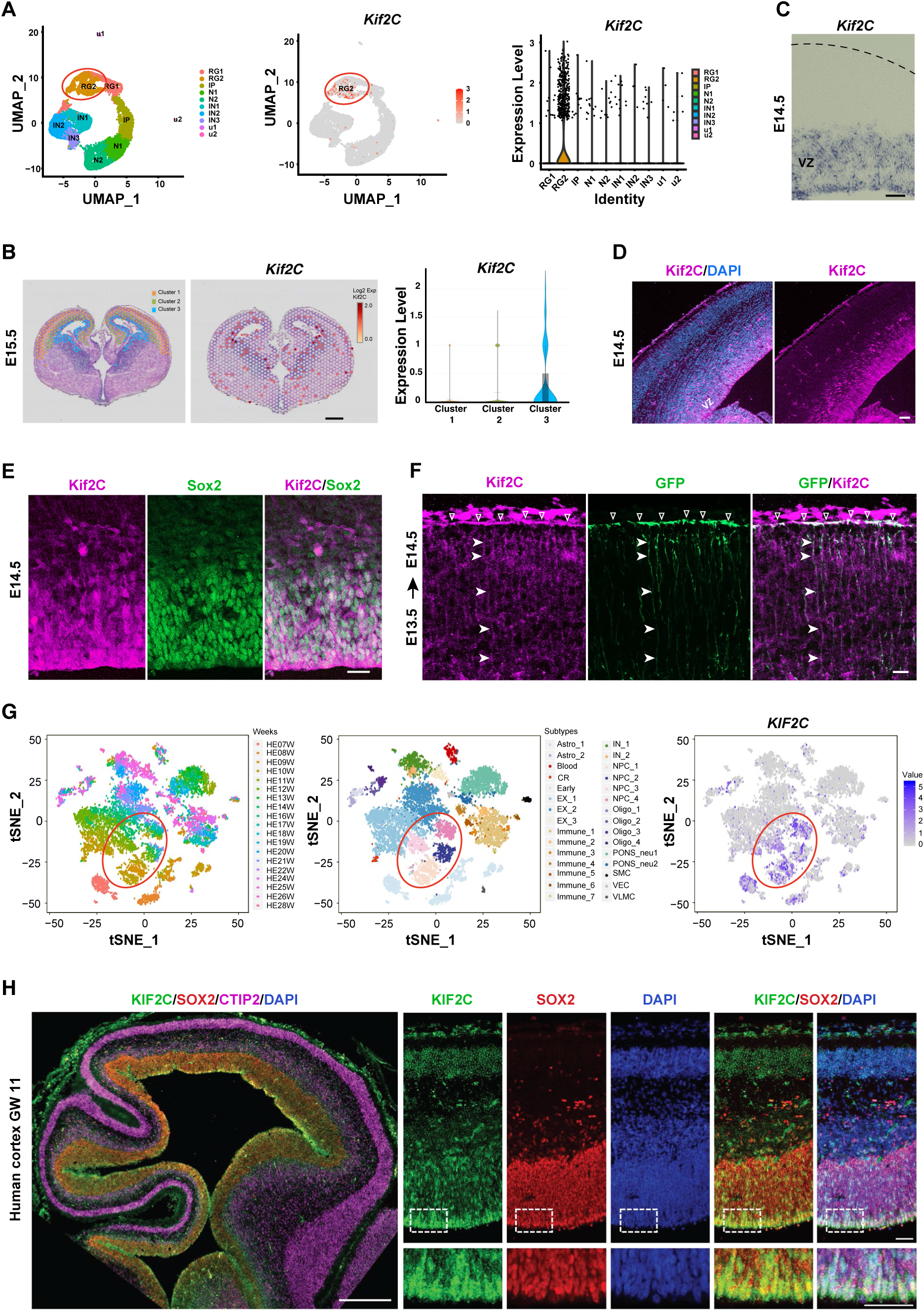
Expression of Kif2C/KIF2C in the developing neocortex. **(A)** Re-analysis of public single-cell RNA-seq dataset of E14.5 mouse cortex (Loo et al., 2019), showing cell type clusters (left), *Kif2C* expression in feature plots (middle), and violin plots of *Kif2C* expression across clusters (right)*. Kif2C* is enriched in the proliferating radial glial cell cluster 2 (RG2). RG1, radial glial cells cluster 1; IP, intermediate progenitor cells cluster; N1 and N2, neuronal clusters 1 and 2; IN1, IN2 and IN3, interneuron clusters 1, 2 and 3. **(B)** Visium spatial gene expression analysis of E15.5 mouse brain, showing spatial gene expression clusters (left), spatial expression of *Kif2C* (middle), and violin plots of *Kif2C* expression across clusters (right). The dataset was derived from Tsai et al., 2024. *Kif2C* is enriched in the cluster 3. Scale bar, 300 μm. **(C)** *In situ* hybridization for *Kif2C* mRNA in the E14.5 mouse cortex. *Kif2C* is localized in the ventricular zone (VZ). Scale bar, 100 μm. **(D)** Immunohistochemistry for Kif2C in the E14.5 mouse cortex. Kif2C is localized in the ventricular zone (VZ). Scale bar, 50 μm. **(E)** Representative images of E14.5 mouse cortex immunostained for Kif2C and Sox2, a marker for radial glial cells. Kif2C is co-localized with Sox2. Scale bar, 20 μm. **(F)** Representative images of E14.5 mouse cortex one day after electroporation of *pCAG-GFP,* immunostained for Kif2C and GFP. White arrowheads indicate Kif2C localization in GFP^+^ radial glial fiber and basal endfeet. Scale bar, 20 μm. **(G)** Analysis of public single-cell RNA-seq dataset from human embryonic and fetal cortex spanning gestational week 7-28 (Fan et al., 2020), showing clusters by gestational week (GW) (left), cell types (middle), and *KIF2C* expression (right). *KIF2C* is enriched in the proliferating cell clusters, including NPC_1, NPC_2, NPC_3, and NPC_4. Early, clusters mainly composed of GW7/8 cortex and GW8/9 pons cells; CC-EX, cortical excitatory neuron; CR, Cajal-Retzius cell; IN, inhibitory neuron; Pons-neu, projection neuron in pons; Oligo, oligodendrocyte; Astro, astrocyte; Immune, immune cells; SMC, smooth muscle cell; VEC, vascular endothelial cell; VLMC, vascular leptomeningeal cell. **(H)** Representative image of GW11 human cortex immunostained for Kif2C and Sox2. Kif2C is co-localized with Sox2 in the ventricular zone (VZ). Boxed areas are magnified in the lower panels. Scale bars, 400 μm (left), 50 μm (top right); 25 μm (bottom right).

We next validated these transcriptomic observations at the mRNA and protein levels. *In situ* hybridization and immunostaining confirmed robust expression of *Kif2C* mRNA (Figure 1C) and Kif2C protein (Figure 1D) in the VZ/subventricular zone (SVZ) of the E14.5 mouse cortex. Co-immunostaining further demonstrated that Kif2C is predominantly expressed in Sox2-positive RGCs (Figure 1E). To define its subcellular localization within RGCs, we labeled RGCs by *in utero* electroporation (IUE) of an EGFP expression vector (*pCAG-EGFP*) at E13.5 and analyzed Kif2C localization at E14.5. Kif2C was detected not only in the soma but also along the basal processes of RGCs, with prominent enrichment at the basal endfeet (Figure 1F). These findings indicate that Kif2C is expressed in proliferative RGCs and is positioned within cellular compartments critical for RGC morphology and cortical scaffold formation.

We then asked whether this expression pattern is conserved in the developing human cortex. Analysis of a publicly available scRNA-seq dataset of human embryonic and fetal cortex spanning gestational weeks (GW) 7–28 [20] showed that *KIF2C* expression is enriched in cortical progenitor populations, with stronger expression at early gestational stages (Figure 1G). To validate this observation at the protein level, we examined human cortical tissue at GW11, corresponding to an early neurogenic stage. KIF2C protein was predominantly detected in the VZ, where RGCs are enriched, and co-immunostaining for Sox2 confirmed its expression in human RGCs (Figure 1H). These data demonstrate that *KIF2C* expression in cortical progenitors is conserved between mouse and human, supporting a conserved role for KIF2C in RGC function during neocortical development.

### Kif2C is required for radial glial scaffold integrity and neuronal migration

To determine the function of Kif2C during corticogenesis, we performed loss-of-function analyses using IUE of a siRNA targeting *Kif2C*. *Kif2C* siRNA or control siRNA was co-electroporated with *pCAG-EGFP* expression vector into the mouse cortex at E14.5, and brains were analyzed at E15.5, E17.5, and E18.5 (Figure 2A). We first confirmed that *Kif2C* siRNA effectively reduced the endogenous Kif2C expression in the embryonic cortex (Figure 2B).

**Figure 2.**
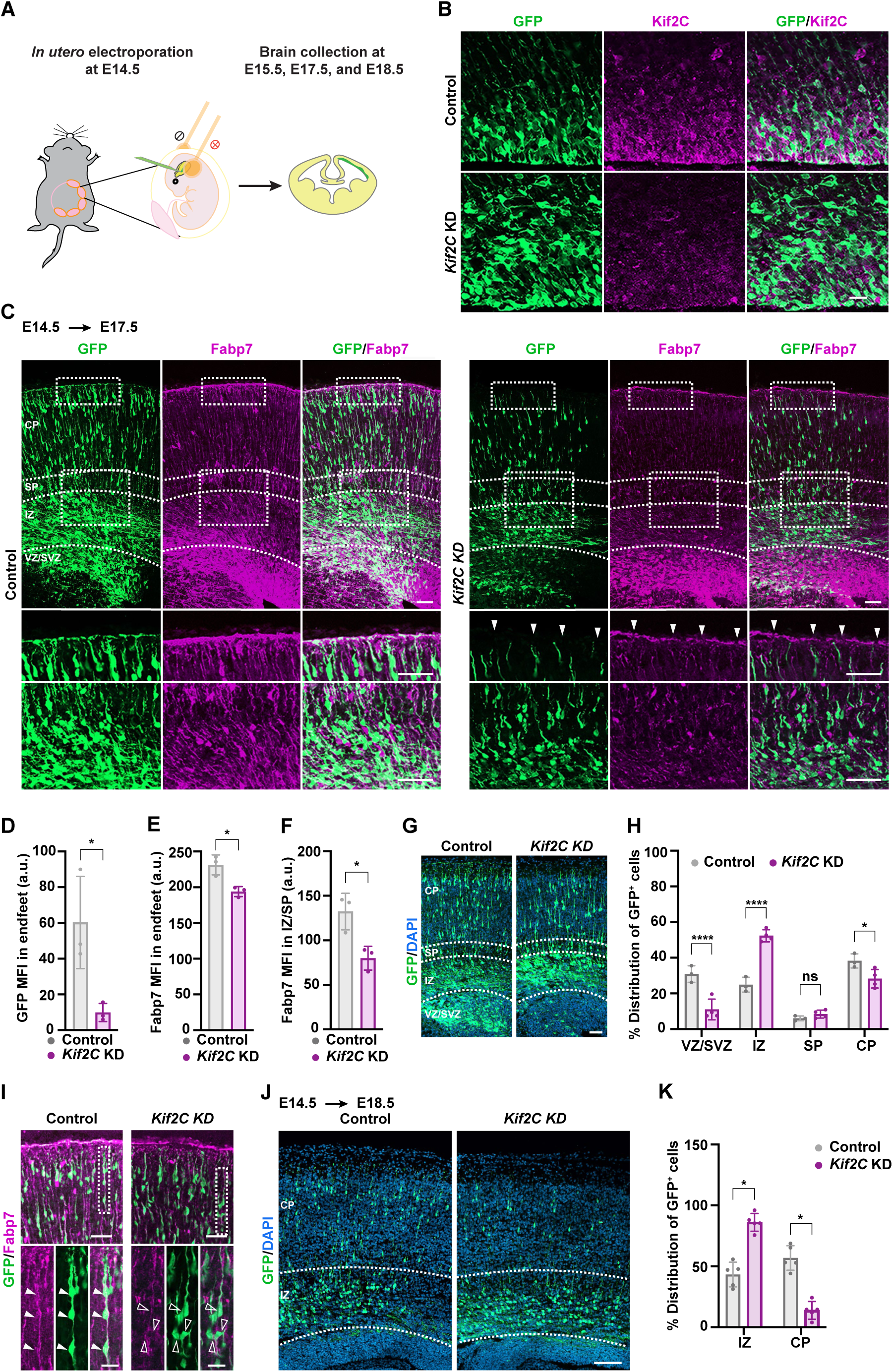
*Kif2C* knockdown disrupts radial glial scaffold and causes neuronal migration defects. **(A)** Schematic diagram of *Kif2C* knockdown (KD) by *in utero* electroporation (IUE). Embryonic mouse cortices were electroporated at E14.5 with control or *Kif2C* siRNA together with *pCAG-EGFP*. **(B)** Representative images of E15.5 mouse cortices immunostained for Kif2C after IUE. Kif2C expression is effectively suppressed by *Kif2C* siRNA. Scale bar, 20 μm. **(C)** Representative image of E17.5 control and *Kif2C-*KD cortices immunostained for GFP and Fabp7. Dashed lines indicate the borders between the ventricular zone/subventricular zone (VZ/SVZ), intermediate zone (IZ), subplate (SP), and cortical plate (CP). Boxed regions are magnified in the lower panels. White arrowheads indicate basal endfeet. Scale bars, 50 μm. **(D-F)** Quantification of mean fluorescence intensity of GFP in basal endfeet **(D)**, Fabp7 in basal endfeet **(E),** and Fabp7 in the IZ/SP region **(F),** as shown in (**C)**. Data are presented as mean ± SD; n = 3 brains per group. Two-tailed Student’s *t*-test, \**p*< 0.05. **(G)** Representative images of E17.5 control and *Kif2C-*KD cortices immunostained for GFP and DAPI. Scale bar, 50 μm. **(H)** Quantification of the distribution of GFP^+^ cells in VZ/SVZ, IZ, SP, and CP, respectively. Data are presented as mean ± SD; n = 3 for control group and n = 4 for *Kif2C*-KD. Two-way ANOVA with Bonferroni’s multiple comparisons test, \**p*< 0.05; \*\*\*\**p*< 0.0001; ns, not significant. **(I)** Representative images of E17.5 control and *Kif2C-*KD cortices immunostained for GFP and Fabp7. Boxed regions are magnified in the lower panels. Closed white closed arrowheads indicate radially migrating neurons aligned with RG fibers; open white arrowheads indicate scattered neurons. Scale bars, 50 μm (top); 20 μm (bottom). **(J)** Representative images of E18.5 control and *Kif2C-*KD cortices immunostained for GFP and DAPI. Scale bar, 100 μm. **(K)** Quantification of the distribution of GFP^+^ cells in the IZ and CP **(K)**. Data are presented as mean ± SD; n = 5 brains per group. Multiple *t*-test, \**p*< 0.05.

We next examined whether Kif2C depletion affects the morphology of radial glial (RG) fibers. To visualize RG fibers, electroporated cortices were co-immunostained for GFP and fatty acid binding protein 7 (Fabp7), a marker of RG fibers [21,22]. At E17.5, GFP^+^ basal endfeet were markedly reduced in *Kif2C* knockdown (KD) cortices compared with controls (Figures 2C and 2D). Consistently, Fabp7-labeled RG fibers in *Kif2C-*KD lost their characteristic elongated organization and instead exhibited discontinuity and disarray (Figure 2C). Quantitative analysis further revealed a significant reduction in Fabp7 signal intensity both at the basal endfeet and within the intermediate zone (IZ)/subplate (SP) region following *Kif2C-*KD (Figures 2E and 2F). Notably, RG scaffold disruption was already detectable at E15.5 (Figure S2A) and became more pronounced by E16.5 (Figure S2B), indicating that Kif2C is required for the maintenance of RG fiber integrity during cortical development.

Because RG fibers provide the scaffold for radial migration of newly generated neurons [23], we next assessed the distribution of GFP^+^ cells across cortical layers. *Kif2C-*KD reduced the proportion of GFP^+^ cells in the VZ and cortical plate (CP), with a corresponding increase in the IZ, compared with controls (Figures 2G and 2H). In control cortices, migrating neurons were closely associated with RG fibers, whereas in the *Kif2C*-KD cortices, GFP^+^ cells appeared scattered and were no longer aligned with the remaining RG fibers (Figure 2I). Furthermore, at E18.5, four days after electroporation, GFP^+^ cells remained abnormally accumulated in the IZ and were correspondingly reduced in number in the CP in *Kif2C*-KD cortices (Figures 2J and 2K). Together, these findings demonstrate that Kif2C is essential for proper RG scaffold organization and radial neuronal migration in the developing cortex.

### Kif2C preserves the radial glial cell pool by maintaining mitotic fidelity

The reduction in GFP^+^ cells within the RGC-enriched VZ of *Kif2C-*KD cortices prompted us to examine whether Kif2C contributes to maintenance of the RGC pool. Two days after IUE, at E16.5, immunostaining for Pax6 (an RGC marker) and Tbr2 (an intermediate progenitor marker) revealed a significant decrease in the proportion of Pax6⁺/GFP⁺ RGCs in *Kif2C*-KD cortices, whereas the Tbr2⁺/GFP⁺ intermediate progenitors was unchanged (Figures 3A–3D). These results indicate that Kif2C depletion selectively reduces the RGC pool.

**Figure 3.**
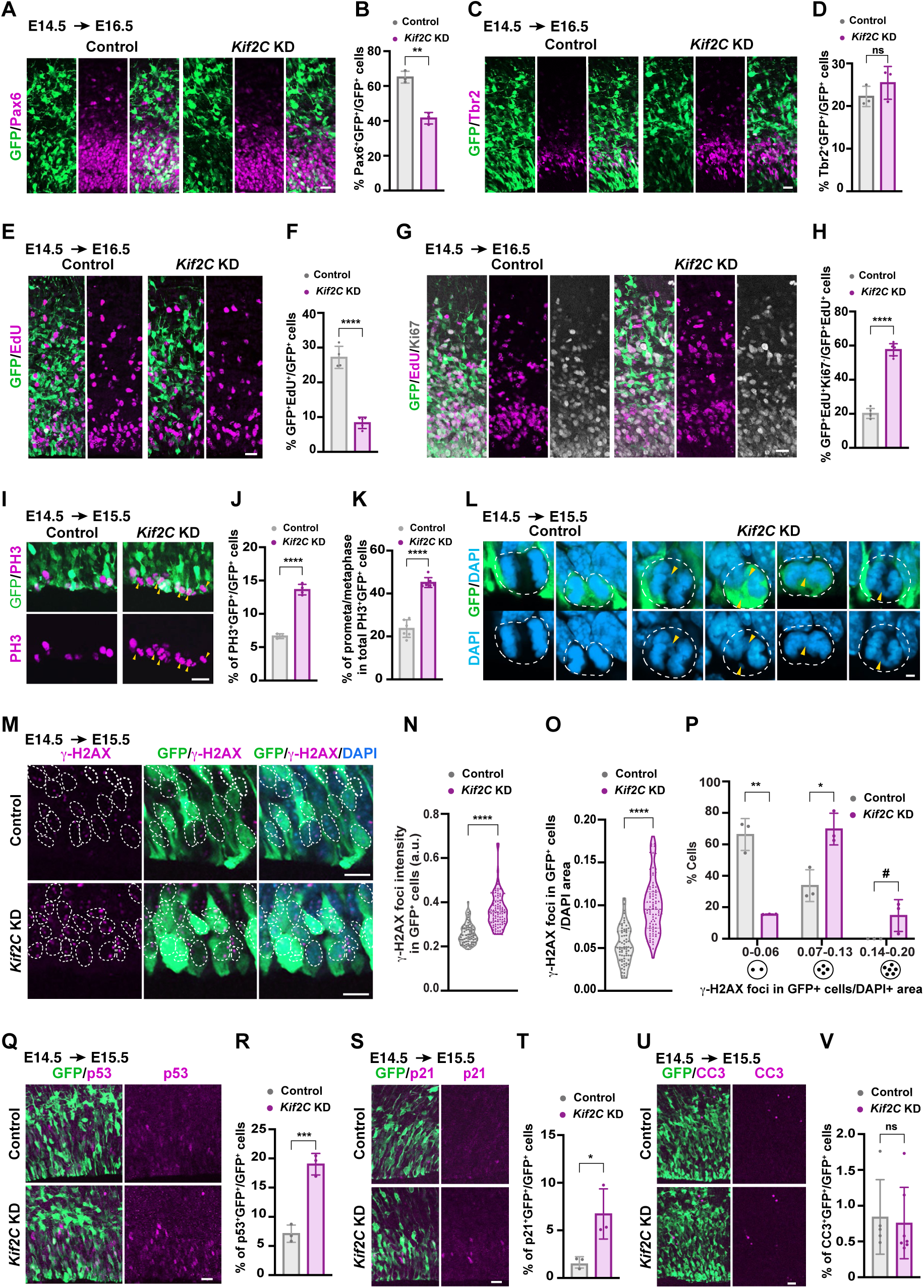
*Kif2C* knockdown impairs RGC maintenance and induces mitotic defects and cell cycle arrest. **(A)** Representative images of E16.5 control and *Kif2C-*KD cortices immunostained for GFP and Pax6. Scale bar, 20 μm. **(B)** Quantification of the percentage of Pax6^+^GFP^+^ cells among total GFP^+^ cells. Data are presented as mean ± SD; n = 3 brains per group. Two-tailed Student’s *t*-test, \**p*< 0.05. **(C)** Representative images of E16.5 control and *Kif2C-*KD cortices immunostained for GFP and Tbr2. Scale bar, 20 μm. **(D)** Quantification of the percentage of Tbr2^+^GFP^+^ cells among total GFP^+^ cells. Data are presented as mean ± SD; n = 3 brains per group. Two-tailed Student’s *t*-test, \**p*< 0.05. **(E)** Representative images of electroporated cortical sections immunostained for GFP and EdU. Scale bar, 20 μm. **(F)** Quantification of the percentage of EdU^+^GFP^+^ cells. Data are presented as mean ± SD; n = 4 for control group and n = 5 for *Kif2C*-KD group. Two-tailed Student’s *t*-test, \*\*\*\**p*< 0.0001. **(G)** Representative images of electroporated cortical sections stained for GFP, Ki67, and EdU. Scale bar, 20 μm. **(H)** Quantification of the percentage of EdU^+^GFP^+^Ki67^−^ cells. Data are presented as mean ± SD; n = 5 for control group and n = 6 for *Kif2C*-KD group. Two-tailed Student’s *t*-test, \*\*\*\**p*< 0.0001. **(I)** Representative images of the E15.5 control and *Kif2C-*KD cortices immunostained for GFP and PH3. Scale bar, 20 μm. **(J, K)** Quantification of the percentage of PH3^+^GFP^+^ cells relative to the total GFP^+^ cells **(J)** and the percentage of prometaphase/metaphase cells relative to the total PH3^+^GFP^+^ cells **(K)**. Data are presented as mean ± SD; n = 4 brains per group for **(J)**; n = 100 cells from 6 samples for control group and n = 176 cells from 7 samples for *Kif2C*-KD group for **(K)**. Two-tailed Student’s *t*-test, \*\*\*\**p*< 0.0001. **(L)** Representative images of E15.5 electroporated cortical sections stained for GFP and DAPI. Yellow arrowheads indicate chromosome bridges in the midzone of anaphase RGCs in *Kif2C-*KD cortices. Scale bar, 2 μm. **(M)** Representative images of E15.5 control and *Kif2C-*KD cortices stained for GFP, γ -H2AX, and DAPI. White scattered lines encircle the boundaries of the nucleus of GFP^+^ cells. Scale bar, 10μm. (**N**-**P**) Quantification of γ-H2AX fluorescence intensity in the GFP^+^ cells (**N**), the number of γ -H2AX foci in the GFP^+^ cells per area of nucleus (**O**), and the percentage of GFP^+^ cells in distinct categories based on the number of ψ-H2AX foci per area of nucleus (**P**). Data are presented as mean ± SD; n = 76 cells from 3 samples for control group and n = 90 cells from 3 samples for *Kif2C*-KD group. Two-tailed Student’s *t*-test, \*\*\*\**p*< 0.0001 (**N, O**); Multiple *t*-tests, \*\**p*< 0.01; \**p*< 0.05. **#**, High γ-H2AX level was only observed in the *Kif2C*-KD cortices (**P**). **(Q)** Representative images of E15.5 control and *Kif2C-*KD cortices immunostained for GFP and p53. Scale bar, 20 μm. **(R)** Quantification of the percentage of p53^+^ cells among GFP^+^ cells. Data are presented as mean ± SD; n = 3 brains per group. Two-tailed Student’s *t*-test, \*\*\**p*< 0.001. **(S)** Representative images of E15.5 control and *Kif2C-*KD cortices immunostained for GFP and p21. Scale bar, 20 μm. **(T)** Quantification of the percentage of p21^+^ cells among GFP^+^ cells. Data are presented as mean ± SD; n = 3 brains per group. Two-tailed Student’s *t*-test, \**p*< 0.05. **(U)** Representative images of E15.5 control and *Kif2C-*KD cortices immunostained for GFP and CC3. Scale bar, 20 μm. **(V)** Quantification of the percentage of CC3^+^ cells among GFP^+^ cells. Data are presented as mean ± SD; n = 3 brains per group. Two-tailed Student’s *t*-test, ns, not significant.

To determine whether this reduction reflects impaired proliferation, we performed IUE at E14.5 and administered EdU two hours before analysis at E16.5 to label S-phase cells. *Kif2C* depletion significantly reduced the proportion of proliferative GFP⁺ cells (Figures 3E and 3F). To assess premature cell cycle exit, we next performed IUE at E14.5, injected EdU at E15.5, and analyzed the cortex at E16.5 using an antibody against Ki67, a marker for proliferating cells [24]. *Kif2C-*KD increased the proportion of EdU^+^/GFP^+^/Ki67^−^ cells, indicating enhanced cell cycle exit (Figures 3G and 3H). Together, these findings demonstrate that Kif2C supports RGC pool maintenance by promoting progenitor proliferation and preventing precocious cell cycle exit.

To explore the mechanism underlying this phenotype, we examined the subcellular localization of Kif2C during mitosis. Kif2C was distributed in both the nucleus and cytoplasm during interphase, accumulated at the centromeres/kinetochores during metaphase, and localized to opposite poles during anaphase and telophase (Figure S3A). Co-staining with α-tubulin, a spindle MT marker, further showed that Kif2C localized to the kinetochore-MT attachment sites at the spindle midplane during metaphase and to spindle poles during anaphase (Figure S3B). These dynamic localization patterns suggest Kif2C regulates kinetochore-MT interactions in mitotic RGCs.

Given the importance of kinetochore-associated proteins in chromosomal congression and segregation [25,26], we next examined mitotic RGCs by phospho-histone H3 (PH3) immunostaining. *Kif2C*-KD cortices showed a significant increase in PH3^+^/GFP^+^ cells compared with control cortices (Figures 3I and 3J), suggesting impaired mitotic progression. Many PH3^+^ cells in *Kif2C*-KD cortices displayed prometaphase/metaphase-like morphology (Figures 3I and 3K), consistent with delayed progression through mitosis [27]. DAPI staining further revealed chromosome bridges during anaphase in *Kif2C*-KD RGCs (Figure 3L), a hallmark of chromosomal missegregation [28]. Together, these findings indicate that Kif2C depletion compromises mitotic fidelity, leading to prometaphase/metaphase accumulation and chromosome segregation defects.

Because mitotic errors can impair genome stability and induce DNA damage [29], and because Kif2C has also been implicated in DNA damage repair [30], we next examined expression of γ-H2AX, a marker of DNA double-strand breaks, in *Kif2C*-KD cortices (Figure 3M). Kif2C- depleted cells exhibited increased γ -H2AX fluorescence intensity and a higher number of γ-H2AX foci per nuclear area compared with controls (Figures 3M–3O). Classification of GFP^+^ cells according to γ -H2AX signal intensity, i.e., low, middle, or high, revealed a shift toward higher γ-H2AX levels in *Kif2C*-KD cortices, with increased proportions of cells in the middle and high categories and a corresponding decrease in the low category (Figure 3P). These findings indicate that Kif2C depletion triggers a DNA damage response in the developing cortex.

To further characterize this response, we examined p53, a tumor suppressor activated by DNA damage [31]. Kif2C downregulation increased p53 expression compared with controls (Figures 3Q and 3R). Because the p53 signaling axis can induce cell cycle arrest or apoptosis [32], we next examined p21, a downstream effector that promotes cell cycle exit [33]. Kif2C depletion increased p21 expression (Figures 3S and 3T). In contrast, immunostaining for cleaved caspase-3 (CC3) did not reveal a significant increase in apoptosis (Figures 3U and 3V). Together, these results indicate that Kif2C depletion activates the γ-H2AX–p53–p21 response, thereby promoting cell cycle exit in RGCs without inducing detectable apoptosis.

### Kif2C knockdown disrupts microtubule organization

Given the critical role of polarized MT networks in maintaining RGC morphology [34], we next examined how *Kif2C-*KD affects MT organization. We first analyzed the MT cytoskeleton in N2a cells, a neuroblastoma-derived model commonly used to assess neurite morphology and MT organization. N2a cells were transfected with *Kif2C* siRNA or control siRNA and stained for α-tubulin. *Kif2C*-KD cells displayed abnormal neurite morphology, including widened neurite shafts and splayed, unevenly organized MT arrays (Figure 4A). Quantification revealed a significant increase in the proportion of cells exhibiting splayed MT within neurite shafts following *Kif2C*-KD (Figure 4B). Neurite length was also significantly reduced in *Kif2C*-KD cells compared with controls (Figure 4C). In addition, Sholl analysis demonstrated increased neurite complexity in *Kif2C-KD* cells, as indicated by a higher number of intersections relative to controls (Figures 4D and 4E). These findings indicate that Kif2C is required for proper organization of the MT cytoskeleton.

**Figure 4.**
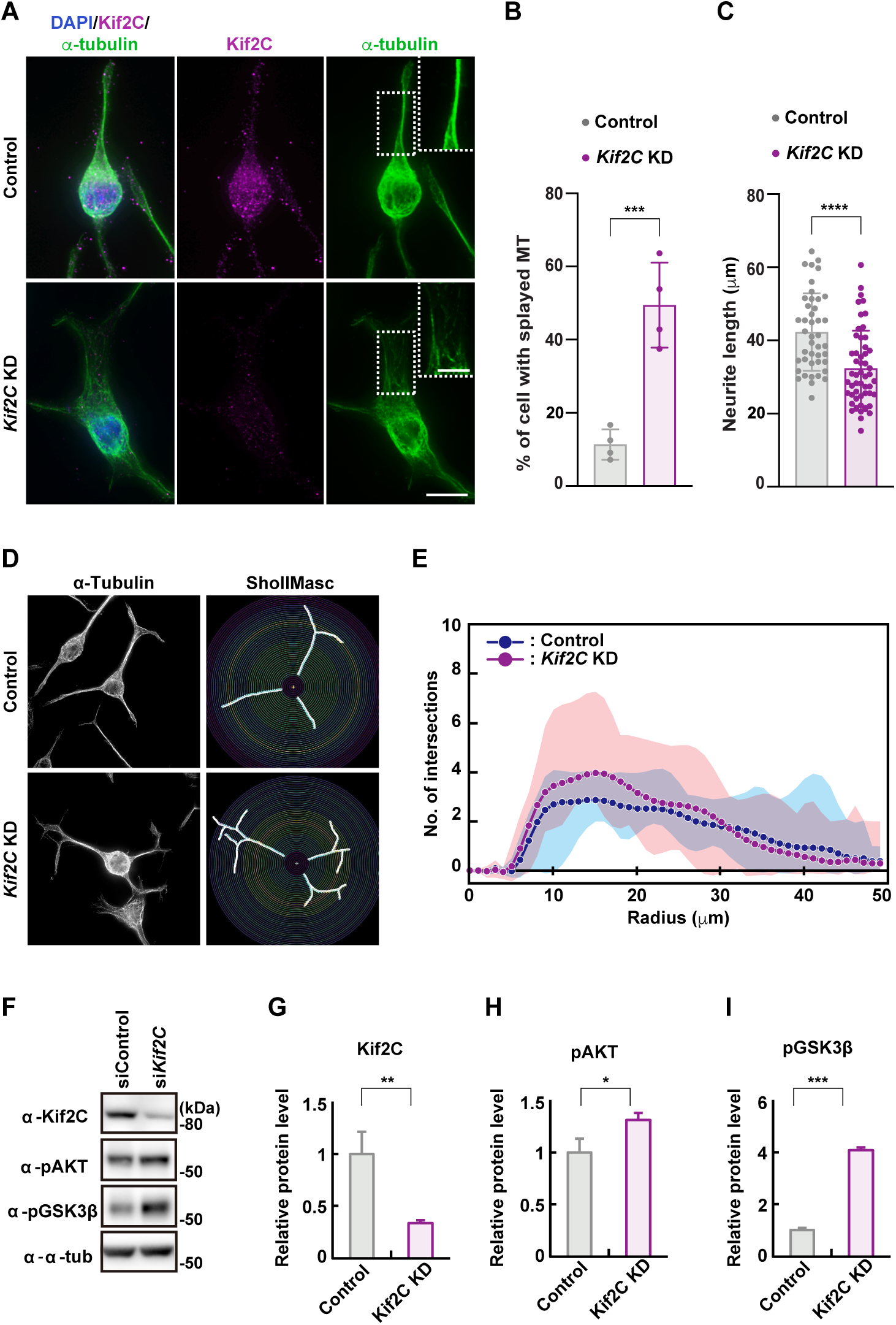
*Kif2C* knockdown impairs microtubule organization and alters AKT-GSK3β signaling. **(A)** Representative images of control and *Kif2C-*KD N2a cells stained for Kif2C, α-tubulin (microtubule marker), and DAPI. Boxed regions are magnified in the insets. Scale bars, 5 μm (inset); 10 μm (bottom). **(B)** Quantification of the percentage of cells with splayed microtubules. Data are presented as mean ± SD; n = 45 cells from 4 replicates for control group and n = 54 cells from 4 replicates for *Kif2C*-KD. Two-tailed Student’s *t*-test, \*\*\**p*< 0.001. **(C)** Quantification of the length of the longest neurite in control and *Kif2C-*KD N2a cells. Data are present as mean ± SD; n = 45 cells from 4 replicates for control group and n = 52 cells from 4 replicates for *Kif2C*-KD. Two-tailed Student’s *t*-test, \*\*\**p*< 0.001. **(D, E)** Sholl analysis of neurite complexity in control and *Kif2C*-KD N2a cells **(D)**. Quantification of intersections per concentric circle **(E)**. Two-way repeated-measures ANOVA, \*\*\*\**p*< 0.001. **(F)** Representative western blots showing KIF2C, phosphorylated AKT (pAKT), and phosphorylated GSK3β (pGSK3β) in lysates from control and *Kif2C*-KD N2a cells. **(G-I)** Quantification of relative protein levels of KIF2C **(G)**, pAKT **(H)**, and pGSK3β **(I)**, normalized to α-tubulin and expressed relative to the control condition, which was set to 1. Data are presented as mean ± SD; n = 3 for control group and n = 3 for *Kif2C*-KD group. Two-tailed Student’s *t*-test, \*\**p*< 0.01 **(G)**, \**p*< 0.05 **(H)**, and \*\*\**p*< 0.001 **(I)**.

We next explored the signaling mechanism that may contribute to this cytoskeletal phenotype. Kif2C has been reported to activate ERK signaling [35], which can negatively regulate AKT, a kinase implicated in MT organization [36,37]. We therefore examined AKT signaling following *Kif2C*-KD. Kif2C depletion significantly increased phosphorylation of AKT at Ser473 compared with controls (Figures 4F–4H). Because GSK3β, a key regulator of MT organization and RG fiber morphology [38,39], acts downstream of AKT, we next measured phosphorylation of GSK3β at Ser9, which represents its inactive form. Consistent with increased AKT activity, *Kif2C*-KD markedly increased inhibitory phosphorylation of GSK3β (Figures 4F and 4I). Together, these results suggest that Kif2C contributes to MT organization, at least in part, by regulating the AKT–GSK3β signaling pathway.

### Kif2C maintains pial basement membrane integrity to prevent neuronal overmigration

Because interactions between RG endfeet and the pial basement membrane establish the boundary between the neocortex and the meninges and provide an essential structural scaffold during corticogenesis [40], we asked whether disruption of RG fibers in *Kif2C-*KD cortices leads to cortical malformations. *Kif2C* depletion caused neuronal ectopias characterized by abnormal neuronal overmigration beyond the marginal zone (Figure 5A, Table S1). To determine whether these ectopias persisted postnatally, we analyzed cortices at postnatal day 7 (P7) by co-immunostaining for GFP and Cux1, a marker of upper layer neurons. Neuronal ectopias were consistently observed in the postnatal *Kif2C-*KD cortices, but not in controls (Figure 5B, Table S1). Such ectopic neuronal accumulation is a characteristic feature of cobblestone lissencephaly in humans [41,42].

**Figure 5.**
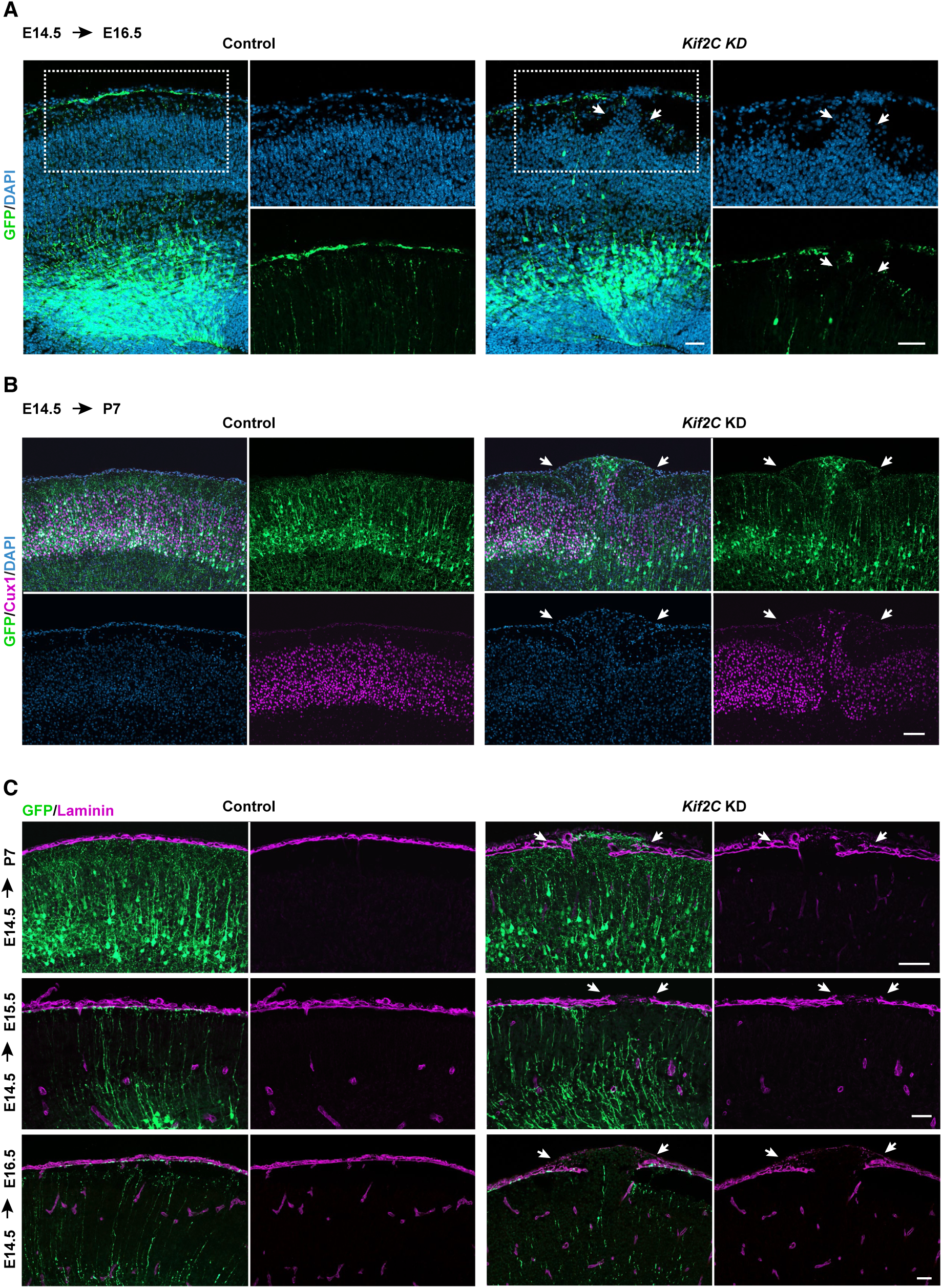
*Kif2C* knockdown results in neuronal ectopia and pial basement membrane breaches. **(A)** Representative images of E16.5 control and *Kif2C-*KD cortices stained for GFP and DAPI. Boxed regions are magnified on the right side. White arrowheads indicate the presence of neuronal ectopia in the *Kif2C-*KD cortex. Scale bars, 50 μm. **(B)** Representative images of P7 control and *Kif2C-*KD cortices stained for GFP, Cux1, and DAPI. White arrowheads indicate the presence of neuronal ectopia in the *Kif2C-*KD cortex. Scale bar, 100 μm. **(C)** Representative images of E15.5, E16.5, and P7 control and *Kif2C-*KD cortices stained for GFP and laminin. White arrowheads indicate pial basement membrane breaches in the *Kif2C-*KD cortices. Scale bar, 100 μm.

The pial basement membrane restricts neuronal positioning and prevents ectopic migration into the extracortical space [43]. We therefore examined its integrity by co-immunostaining for GFP and laminin, a major basement membrane component. In control cortices, laminin signal was continuous and uniform along the pial surface, whereas *Kif2C*-KD cortices displayed focal laminin discontinuities that coincided with sites of neuronal overmigration at P7 (Figure 5C), indicating that pial basement membrane disruption contributes to ectopic phenotype. To define the temporal progression of this defect, we performed a time-course analysis. Disruption of the pial basement membrane was already detectable at E15.5, one day after electroporation, and became more pronounced at E16.5, two days after electroporation, in *Kif2C*-KD cortices (Figure 5C). Together, these findings demonstrate that Kif2C is required to maintain pial basement membrane integrity, thereby preventing neuronal overmigration and a cobblestone-like cortical malformation.

### A patient-derived truncating *KIF2C* variant fails to rescue Kif2C loss-of-function phenotypes

To examine the clinical relevance of our findings, we analyzed previously unreported *KIF2C* variants identified by whole-exome sequencing in two unrelated individuals with severe neurodevelopmental abnormalities: one from a Pakistani family and the other from a Polish family (Figures 6A, and 6B; Table S2). The Pakistani individual carried a homozygous nonsense variant in *KIF2C* (NM_006845: c.690G>A; p.Trp230*), whereas the Polish individual carried a distinct *de novo* frameshift variant (NM_006845.4:c.1471dup, NP_006836.2:p.Val491GlyfsTer7). Despite differences in inheritance pattern and variant position, both individuals showed overlapping neurodevelopmental features, including microcephaly, intellectual disability, global developmental delay, motor impairment, abnormal muscle tone or movement, and severely impaired language development (Table S2). These shared clinical features consistently suggest an essential role for KIF2C in human brain development.

**Figure 6.**
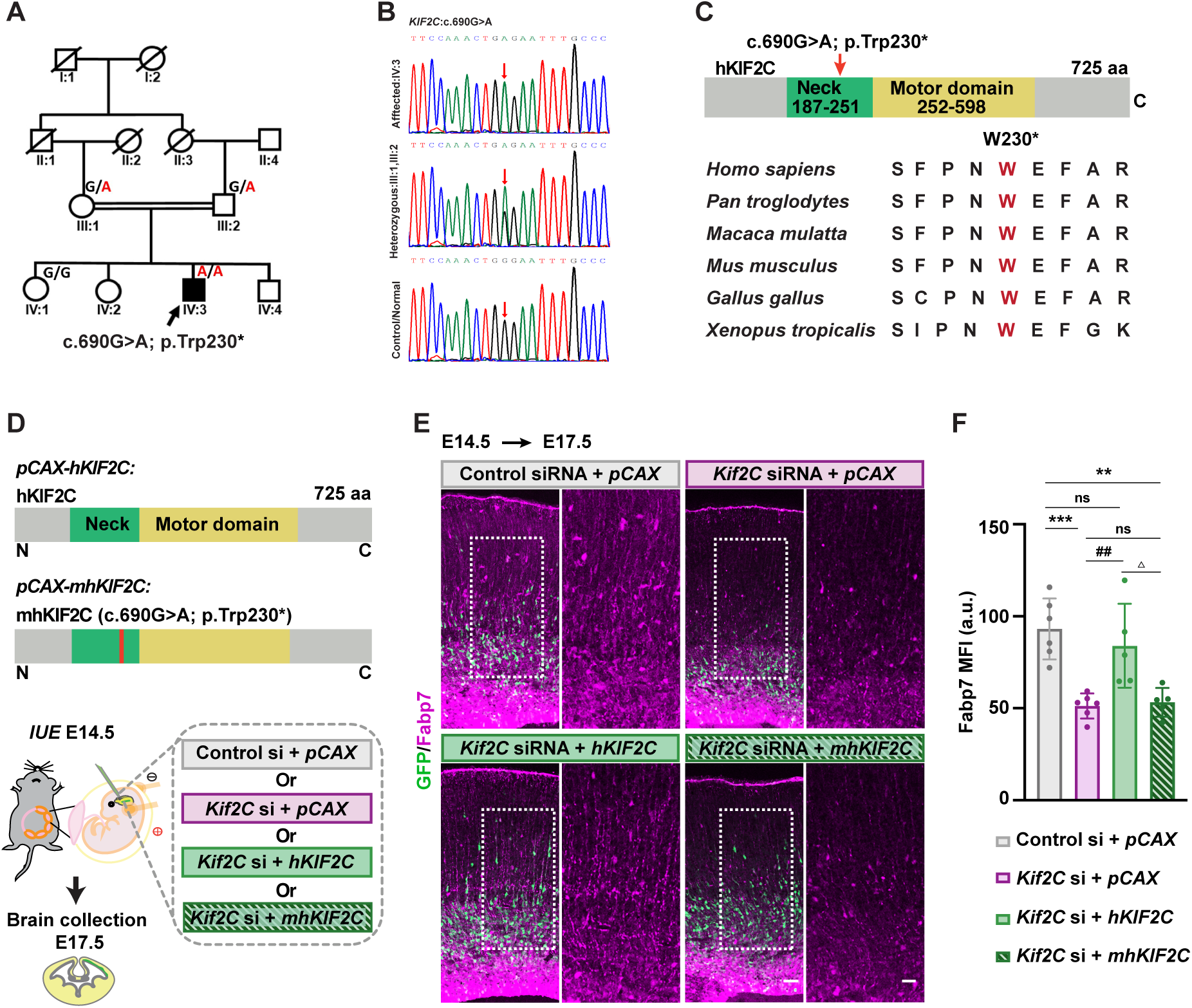
*Kif2C* knockdown phenotypes are rescued by WT human KIF2C but not by a patient-derived variant. **(A)** Pedigree of a patient with a neurodevelopmental disorder carrying a homozygous *KIF2C* nonsense variant, c.690G>A (p.Trp230*). **(B)** Electropherograms confirming the c.690G>A substitution, indicated by red arrows, in the affected individual (homozygous), parents (heterozygous), and a healthy control (wild-type). **(C)** Schematic diagram of KIF2C protein domains showing the position of the patient-derived p.Trp230* variant (top). Amino acid conservation of the mutated region across species is shown below. **(D)** Schematic diagram of WT human *KIF2C* and mutant *KIF2C* constructs (top), and overview of the rescue experiment (bottom). **(E)** Representative images of E17.5 mouse cortices stained for GFP and Fabp7 (E). Boxed regions indicate areas shown at higher magnification. Scale bars, 50 μm (left); 25 μm (right). **(F)** Quantification of Fabp7 mean fluorescence intensity. Data are presented as mean ± SD; n = 6 brains for control siRNA + *pCAX* group; n = 6 brains for *Kif2C* siRNA + *pCAX* group; n = 5 brains for *Kif2C* siRNA + *pCAX-hKIF2C* group; n = 5 brains for *Kif2C* siRNA + *pCAX-mhKIF2C*. One-way ANOVA with Tukey’s multiple comparisons test. *p* values: \*\**p*< 0.01, \*\*\**p*< 0.001, ns: not significant (vs control siRNA + *pCAX* group); ^##^*p*< 0.01, ns: not significant (vs *Kif2C* siRNA + *pCAX* group); ^Δ^*p*< 0.05 (vs *Kif2C* siRNA + *pCAX-mhKIF2C* group).

The p.Trp230* variant identified in the Pakistani individual affects a tryptophan residue that is highly conserved across vertebrates and introduces a premature stop codon upstream of the motor domain, predicting a severe loss of KIF2C protein function (Figure 6C). The Polish p.Val491GlyfsTer7 variant affects the motor domain and was associated with additional neurological and sensory features, including profound motor delay, congenital sensorineural deafness, abnormal visual responses, and brain stem hypoplasia involving the lower pons, medulla oblongata, and pontomedullary junction (Figures S4A and S4B; Table S2). Thus, although the two clinical presentations were not identical, both damaging variants support a link between impaired KIF2C function and human neurodevelopmental pathology. Given that the Pakistani individual carried a homozygous truncating variant predicted to cause pronounced loss of protein function, and because our mechanistic analyses focused on cortical development, we prioritized the p.Trp230* allele for functional validation.

To test whether the p.Trp230* variant retains KIF2C activity in the developing cortex, we performed rescue experiments in the mouse *Kif2C*-KD model (Figure 6D). Wild-type human *KIF2C* (*WT hKIF2C*) was first validated by IUE, which confirmed robust protein expression in the embryonic cortex (Figure S5A). Co-electroporation of *WT hKIF2C* with mouse *Kif2C* siRNA at E14.5 restored KIF2C protein levels, indicating that the human construct was resistant to the siRNA targeting mouse Kif2C (Figures S5B, S6A, and S6B). We then assessed whether *WT hKIF2C* or the p.Trp230* mutant construct (*mhKIF2C*) could rescue the RG fiber defects caused by mouse *Kif2C*-KD. Co-electroporation of *WT hKIF2C* with *Kif2C* siRNA restored Fabp7^+^ RG fiber organization at E17.5 to a level comparable to that observed in controls (Figures 6E and 6F). In contrast, the p.Trp230* mutant (*mhKIF2C*) failed to rescue the disrupted RG scaffold, and Fabp7 signal intensity remained significantly reduced, similar to that observed in *Kif2C*-KD alone (Figures 6E and 6F). Together, these findings indicate that the patient-derived p.Trp230* variant behaves as a loss-of-function allele *in vivo* and support the idea that impaired KIF2C function contributes to human neurodevelopmental pathology.

## Discussion

In this study, we identify the MT-depolymerizing kinesin Kif2C as a critical regulator of RGC architecture and mitotic progression during cortical development. Kif2C is enriched in RGCs and localizes to both RG fibers and mitotic structures, supporting dual roles in cytoskeletal organization and cell division. Loss of Kif2C disrupts RG scaffolds, impairs neuronal migration, reduces the RGC pool through premature cell cycle exit, and ultimately causes cobblestone-like cortical malformations. These findings suggest that Kif2C coordinates MT dynamics and progenitor behavior to ensure proper neocortical development.

RGCs serve both as neural progenitors and as scaffolds for radial neuronal migration through their elongated basal fibers [44]. In contrast to the kinesin-13 family member Kif2A, which is predominantly expressed and functions in neurons [45,46], we found that Kif2C is prominently expressed in RGCs and localizes along RG fibers. Kif2C depletion disrupted RG fiber organization, producing shortened and discontinuous fibers with bead-like swellings. This structural disruption was accompanied by severe impairment of neuronal migration. Given the dependence of radial migration on an intact RG scaffold, these findings suggest that Kif2C is required for neuronal positioning primarily through maintenance of RG fiber integrity.

The morphology of RG fibers is largely determined by MT organization rather than actin microfilaments [47,48]. MT-associated proteins regulate the formation and maintenance of polarized MT networks in RGCs [49,50], and our findings suggest that Kif2C contributes to this process by maintaining MT organization within RG fibers. Kif2C deficiency disrupted MT alignment and RG fiber morphology and was accompanied by increased AKT phosphorylation and inhibitory phosphorylation of GSK3β. Because GSK3β is a well-established regulator of MT dynamics, and its inhibitory phosphorylation can influence MT stability and bundling [51,52], these findings support a model in which Kif2C loss leads to sustained AKT activation, GSK3β inhibition, and consequent MT disorganization. Previous studies have shown that MT depolymerization can reduce AKT activity [53], raising the possibility that loss of Kif2C-mediated MT depolymerizing activity perturbs feedback regulation of this pathway. Although Kif2C has also been implicated in ERK signaling in other cellular contexts [35], whether this pathway contributes to Kif2C-dependent regulation of RGC morphology remains to be determined. Together, these findings suggest that Kif2C maintains RG fiber integrity through coordinated regulation of MT organization and intracellular signaling.

In addition to its localization along RG fibers, Kif2C was detected at centromeres/kinetochores in mitotic RGCs, suggesting that it supports both radial scaffold integrity and mitotic fidelity in cortical progenitors. MT depolymerizing proteins are essential for correcting improper kinetochore-MT attachments and ensuring accurate chromosome movement during mitosis [54,55]. Accordingly, the increase in mitotic abnormalities and chromosome segregation errors after Kif2C depletion indicates that Kif2C is required for proper kinetochore-MT dynamics and mitotic progression in RGCs. This function is particularly relevant to cortical growth, because depletion of neural progenitors is a major cellular mechanism underlying microcephaly [56,57]. Consistent with this concept, mitotic defects in neural progenitors can alter cell fate, promote cell cycle exit, and compromise cortical development [58–60]. Moreover, mitotic errors in RGCs can trigger DNA damage responses [61], and Kif2C deficiency was accompanied by increased γ-H2AX and p53 levels without a marked increase in apoptosis. Instead, the associated elevation of p21 suggests that Kif2C-deficient progenitors preferentially undergo premature cell cycle exit, leading to depletion of the RGC pool. Thus, beyond its role in RG morphology, Kif2C is essential for preserving the progenitor population required for normal cortical expansion.

Loss of Kif2C also caused disruption of the pial basement membrane, accompanied by neuronal overmigration and ectopic neuronal clusters. Because the pial basement membrane is maintained through interactions between RG endfeet and the meningeal surface [62,63], these findings suggest that Kif2C is required to preserve the structural integrity of the pial boundary during corticogenesis. The RG fiber defects observed after Kif2C depletion further support the idea that abnormal RG morphology compromises pial surface organization. Consistent with this possibility, Kif2C has been implicated in focal adhesion turnover in other cell types [15], raising the possibility that Kif2C contributes to adhesion-dependent stabilization of RG endfeet at the pial surface. Disruption of this process may weaken basement membrane integrity and permit neuronal overmigration. Notably, a recent study using Fabp7-deficient human fetal brain slice cultures demonstrated that disruption of RG scaffold integrity is accompanied by neuronal mispositioning and transcriptional changes in cytoskeletal regulatory pathways [64], further highlighting the importance of cytoskeletal programs in maintaining RG architecture during human cortical development.

Our findings further support the relevance of KIF2C-dependent mechanisms to human brain development. Consistent with our mouse data, KIF2C is robustly expressed in human RGCs. We identified two damaging *KIF2C* variants in unrelated individuals with overlapping neurodevelopmental phenotypes, including microcephaly, intellectual disability, global developmental delay, and motor impairment. Functional rescue experiments showed that, unlike wild-type human *KIF2C*, the patient-derived p.Trp230* variant failed to restore RG fiber integrity in the mouse *Kif2C*-KD cortex. These findings provide functional evidence that the p.Trp230* variant behaves as a loss-of-function allele *in vivo* and support a role for impaired KIF2C function in the pathogenesis of human cortical developmental abnormalities.

The second individual, carrying a distinct *de novo* heterozygous *KIF2C* frameshift variant, also presented with severe neurodevelopmental impairment and microcephaly, together with additional neurological and sensory features and hindbrain abnormalities on MRI. Although this variant was not functionally modeled in the present study, this case broadens the potential clinical spectrum associated with damaging *KIF2C* variants. In particular, the presence of brain stem hypoplasia raises the possibility that consequences of KIF2C dysfunction may extend beyond the cerebral cortex. This interpretation is biologically plausible, as public expression data from the developing cerebellum indicates *Kif2C* specific expression in cerebellar progenitors (Figure S4C) [65], and Purkinje cell-specific deletion of *Kif2C* in mice has been reported to impair synaptic transmission and motor coordination [66]. Thus, while our experimental data establish a mechanistic link between Kif2C loss and cortical developmental abnormalities, the human cases suggest that KIF2C dysfunction may contribute to a broader neurodevelopmental spectrum.

Together, our findings identify Kif2C as a key regulator of RGC integrity that links MT dynamics, mitotic progression, and cortical architecture during development. By connecting defects in cytoskeletal organization and cell division to impaired radial scaffold function, this work provides a framework for understanding how perturbations in progenitor biology give rise to cortical malformations. More broadly, these findings suggest that coordinated regulation of cytoskeletal dynamics and mitotic fidelity is essential for maintaining tissue architecture in the developing brain.

### Limitations of the study

While this study establishes Kif2C as an important regulator of RG integrity and cortical development, several points warrant further investigation. First, our analysis of the p-AKT–GSK3β pathway was conducted primarily *in vitro*. This reflects the technical difficulty of quantifying subtle signaling changes in the developing cortex, where IUE targets only a limited subset of RGCs, and currently available reagents are not optimal for sensitive *in vivo* biochemical analysis. Nevertheless, the *in vitro* findings provide a useful mechanistic framework for understanding how Kif2C loss may affect MT organization and RG fiber morphology. Second, although we identified human *KIF2C* variants and confirmed KIF2C expression in the developing human cortex, functional testing was performed in the mouse embryonic cortex. Given species differences in cortical architecture, future studies using human cortical organoids or gyrencephalic model systems will help further define the relevance of these mechanisms to human cortical development.

## Data Availability

This study includes no data deposited in external repositories.

## Acknowledgments

We thank Drs. Satoshi Miyashita, Mikio Hoshino, and Yoshio Wakamatsu for their technical advice and valuable comments. We are grateful to Ms. Sayaka Makino for animal care and technical support, and to all members of the Osumi laboratory for helpful discussions. This work was supported by JSPS KAKENHI grants (#JP23K06297, #JP26K09819) to T.K and by JSPS KAKENHI grant (#JP24K02203) and AMED grant (#JP21wm0425003) to N.O. Part of this study was supported by the Support System for Young Researchers to use research equipment, instruments, and devices at Tohoku University. For the Polish case, clinical evaluation and data acquisition were supported by a “Schlüsselprojekt” grant from the Else Kröner-Fresenius-Stiftung (2022_EKSE.185) to M.Z.; by funding from the EJP RD Joint Transnational Call 2022; and by the German Federal Ministry of Education and Research, BMBF, Bonn, Germany, through the project PreDYT “PREdictive biomarkers in DYsTonia,” 01GM2302. M.Z. also received support from the BMBF and the Free State of Bavaria under the Excellence Strategy of the Federal Government and the Länder, as well as the Technical University of Munich - Institute for Advanced Study.

## Author Contributions

Conceptualization: S.N., T.K., and N.O. Methodology and Investigation: S.N. and T.K. performed and analyzed the in vivo knockdown experiments; K.I. performed the in vitro N2a cell experiments; H.Y.C. performed the human cortical slice experiments; S.N. generated the mutant human KIF2C plasmid; J.W.T. performed the Visium spatial transcriptomics analysis; and A.K., M.K., M.M.B., M.Z., and M.H.T. contributed clinical data from human patients. Writing – Original Draft: S.N., T.K., and N.O. Writing – Review & Editing: S.N., T.K., K.I., H.Y.C., S.N., A.K., M.K., M.H.T., M.M.B., J.W.T., L.N., K.T., and N.O. Supervision: T.K. and N.O. Funding Acquisition: T.K., M.Z., and N.O.

## Material and Methods

### Identification of *Kif2C* expression using scRNA-seq data

The scRNA-seq datasets analyzed in this study were obtained from publicly available databases: mouse cortex (GSE123335) [18], human cortex (GSE120046) [20], and mouse cerebellum (GSE178546) [65]. Data normalization, dimensionality reduction, clustering, and visualization of scRNA-seq data were performed using Seurat (v4.3) [67]. The Seurat functions NormalizeData, RunPCA, RunUMAP, FindNeighbors, and FindClusters were used with default parameters.

### Animals

C57BL/6J mice purchased from CLEA Japan were used as wild-type (WT) mice. Embryonic day 0.5 (E0.5) was defined as midday on the day when a vaginal plug was detected. All animal experiments were conducted in accordance with the National Institutes of Health guidelines for the care and use of laboratory animals. Experimental procedures were approved by the Ethics Committee for Animal Experiments of Tohoku University Graduate School of Medicine (approval number: 2019 MdA-018-07).

### *In situ* hybridization

Dorsal telencephalon tissue from WT E14.5 mice was used for RNA extraction with the RNeasy Plus Mini Kit (QIAGEN). cDNA was synthesized from the extracted RNA using the Superscript™ III First-Strand Synthesis System for RT-PCR (Invitrogen). To generate the RNA probe, a cDNA fragment encoding *Kif2C* (NM_134471, nucleotides 134-643) was cloned into pBluescript II SK (–) (Stratagene). Digoxigenin (DIG)-labeled RNA probes were synthesized using the DIG RNA Labeling Kit (Roche). *In situ* hybridization was performed as described previously [68]. Images were acquired using the BZ-X710 fluorescence microscope system (KEYENCE).

### Immunohistochemistry

Immunohistochemistry was performed as described previously [68]. Briefly, frozen brain sections were incubated with primary antibodies diluted with 3% bovine serum albumin in Tris-buffered saline with 0.1% Triton X-100. The following primary antibodies were used: chicken anti-GFP (1:1000; Abcam), rabbit anti-Kif2C (1:1000; Proteintech Group), rabbit anti-Pax6 (1:1000; MBL), rabbit anti-Tbr2 (1:1000; Abcam), rabbit anti-Ki67 (1:500; Abcam), rabbit anti-p53 (1:1000; Leica Biosystems), rabbit anti-active caspase-3 (1:1000; BD Biosciences), rabbit anti-FABP7 (1:1000; Abcam), rabbit anti-Cux1 (1:1000; Santa Cruz Biotechnology), rat anti-Tbr2 (1:1000; Invitrogen), mouse anti-p21 (1:1000; Proteintech Group), mouse anti-phospho-histone H3 (1:1000; Cell Signaling Technology), mouse anti-α-tubulin (1:200, Sigma), and mouse anti-γ-H2AX (1:1000; Millipore). The following secondary antibodies were used: Cy3-conjugated affinity purified anti-rabbit or anti-mouse IgG donkey antibodies (1:500; Jackson Immunoresearch Laboratories), and Alexa 488-conjugated affinity-purified anti-mouse IgG or anti-chicken IgY goat antibodies (1:500; Jackson Immunoresearch Laboratories). Nuclei were counterstained with 4′,6-diamidino-2-phenylindole dihydrochloride (DAPI) (1:1000; Sigma). Images were acquired using a Zeiss LSM800 confocal laser scanning microscope (Carl Zeiss).

### *In utero* electroporation

*In utero* electroporation (IUE) was performed with slight modification to a previously described protocol [69]. Pregnant WT mice at E14.5 were anesthetized with isoflurane. A mixture of siRNA and pCAG-EGFP was injected into the lateral ventricle of the brain using a glass capillary and electric microinjector system (IM-300, Narishige). Five electrical pulses of 40 V, each with a duration of 50 ms and delivered at 1-s intervals, were applied across the uterine wall using forceps-type electrodes (LF650P5, BEX) connected to an electroporator (CUY-21, BEX). The final concentrations of siRNA and *pCAG-EGFP* were 2 μg/μl and 0.5 μg/μl, respectively. Stealth RNAi against mouse *Kif2C* was purchased from Invitrogen. The sequences were as follows:

1. *Kif2C* RNAi (5′- CCGAGATGACAGATCAGCCAGACTA-3′)
2. Scrambled control RNAi (5′- CCGAGTAAGACGACTGACCAGACTA-3′).

### EdU labeling

5-Ethynyl-2’-deoxyuridine (EdU) labeling was performed as described previously [70]. EdU was administered intraperitoneally to pregnant mice at 50 mg/kg. For cell proliferation analysis, pregnant mice injected with EdU at E16.5 were sacrificed 2 h after injection. For cell cycle exit analysis, pregnant mice were injected with EdU at E15.5 and sacrificed 24 h later at E16.5. EdU detection was performed using the Click-iT^TM^ Plus EdU Alexa Fluor^TM^ 555 Imaging Kit (Invitrogen) according to the manufacturer’s protocol.

### N2a cell culture and transfection

N2a cells (mouse neuroblastoma cells, TKG 0509; Institute of Development, Aging and Cancer, Tohoku University, Sendai, Miyagi, Japan) were cultured on glass coverslips. Cells were fixed with 3% paraformaldehyde in PBS containing 137 mM NaCl, 2.7 mM KCl, 10 mM Na2HPO4, and 1.8 mM KH2PO4, pH 7.4, for 10 min at 37°C, and then permeabilized with 1% Triton X-100 in PBS for 5 min. RNAi duplexes were transfected into cells at a final concentration of 50 nM using Lipofectamine RNAiMAX reagent (Thermo Fisher Scientific) 48 h before experiments.

### Immunocytochemistry of cultured cells

Fixed N2a cells were incubated with primary antibodies diluted at 1:2000 for 1 h, washed with PBS containing 0.02% Triton X-100, and incubated for 1 h with Alexa Fluor–488/594-conjugated secondary antibodies (Thermo Fisher Scientific; A11029/A11032 for mouse IgG, A11034/A11037; 1:3,000). All antibody incubations were performed in PBS containing 0.02% Triton X-100. After final washes, cells were mounted with ProLong Glass Antifade Mountant (Thermo Fisher Scientific). Z-image stacks were acquired at 0.2-µm intervals using an Olympus IX-71 inverted microscope controlled by DeltaVision softWoRx (Cytiva), equipped with a 60x/1.42 NA Plan Apochromat oil objective lens (Olympus) and a CoolSnapHQ2 charge-coupled device camera (Photometrics). When necessary, images were deconvolved using an enhanced ratio algorithm, medium noise filtering and 10 iterations per channel. Image stacks were projected and saved as TIFF files and Photoshop files.

### Western blotting

N2a cells were lysed in TNE-N buffer containing 1% NP-40, 100 mM NaCl, 10 mM Tris-HCl, pH 7.5, and 1 mM EDTA, supplemented with cOmplete Protease Inhibitor Cocktail (Merck) and Phosphatase Inhibitor Cocktail (Nakarai Tesque). Cell lysates were boiled for 10 min in 4× NuPAGE LDS sample buffer (Thermo Fisher Scientific). Proteins were separated using the NuPAGE SDS gel system (Thermo Fisher Scientific), transferred onto polyvinylidene difluoride membranes (Amersham Hybond-P; Cytiva), and subjected to immunodetection. The following primary antibodies were used: rabbit anti-Kif2C (1:2000; Proteintech Group), rabbit anti-phospho-Akt (Ser473, D9E) (1:2000; Cell Signaling), rabbit anti-phospho-GSK-3 beta (Ser9, 5B3) (1:2000; Cell Signaling) and mouse anti-α-tubulin B-5-1-2 (1:2000, Merck). Blocking and antibody incubations were performed in 3% nonfat dry milk. Proteins were visualized using horseradish peroxidase-conjugated secondary antibodies (1:10,000; SouthernBiotech) and ECL Prime Western Blotting Detection Reagents (Cytiva) according to the manufacturer’s instructions.

### Human subjects

All human studies were conducted in accordance with the Declaration of Helsinki. Written informed consent was obtained from the legal guardians of the affected individuals. This study includes two affected individuals from two unrelated families.

For the Pakistani case, a consanguineous family with one affected male individual from Khyber Pakhtunkhwa Province, Pakistan, was recruited. The study was approved by the Institutional Review Board (DLSBBC:2022-04). Clinical evaluation was performed by a local clinical neurologist at District Headquarters Hospital, Bannu, Khyber Pakhtunkhwa. Relevant clinical and physical information, including age, sex, family history, and consanguinity, was recorded.

For the Polish case, one affected female individual was investigated as part of an ethics-approved study on dystonia genetics at the Technical University of Munich. The study was approved by the Bioethics Committee for Scientific Study of Medical University of Gdańsk, Poland (approval number: KB/547/2025). The patient was diagnosed at a specialized tertiary pediatric neurology ward in Northern Poland, the Department of Developmental Neurology at the Medical University of Gdansk. Clinical assessments included retrospective analysis of medical records, a detailed medical interview, and prospective neurological examinations with video recording. Clinical evaluations were performed by certified pediatric neurologists and medical geneticists. Brain MRI images were evaluated independently by two certified radiologists.

### Human genomic DNA extraction and purification

Genomic DNA was isolated from peripheral blood samples obtained from the affected individuals and both biological parents for two independent families.

For the Pakistani case, genomic DNA was extracted from peripheral blood using the QIAquick DNA Extraction Kit (Qiagen, Hilden, Germany). DNA concentrations were determined using a Nanodrop-2000 spectrophotometer (Thermo Fisher Scientific, Waltham, MA, USA).

For the Polish case, genomic DNA was isolated from peripheral blood leukocytes according to the standard methods as previously described [71].

### Whole exome sequencing study

To identify the underlying genetic etiology, trio-based whole-exome sequencing was performed using two distinct platforms.

For the Pakistani case, libraries were prepared using the Ion AmpliSeq Library Kit 2.0 (Thermo Fisher Scientific, Waltham, MA, USA). Libraries were diluted at 100 pmol/L and subjected to emulsion PCR using the OneTouch 2 instrument with an Ion Proton I Template OT2 200 Kit v3. Template-positive Ion Sphere Particles were enriched using the Ion One Touch Enrichment System (Thermo Fisher Scientific). The Ion Proton I chip was prepared and loaded on the Ion Proton sequencer. Sequence data were aligned to the human reference genome GRCh37/hg19 using the Ion Torrent Mapping Alignment Program (ITMAP, Thermo Fisher Scientific). Multi-allelic substitutions and indels were genotyped using the Torrent Variant Caller plugin (Thermo Fisher Scientific). Variants were annotated using ANNOVAR (http://www.wannovar.usc.edu/), and sequence reads were visualized using the Integrated Genomic Viewer (IGV, http://www.broadinstitute.org/igv/). Variants were screened according to genomic location, population frequency, and variant type. Variants were filtered using a minor allele frequency cutoff of 1% in the Exome Variant Server (http://evs.gs.washington.edu/EVS/), GnomAD (https://gnomad.broadinstitute.org), and the 1000 Genomes Project databases (http://www.1000genomes.org/). The analysis focused on non-synonymous single-nucleotide variants, including missense, nonsense, splice-site, and frameshift variants. Functional effects were predicted using Polyphen-2 (http://genetics.bwh.harvard.edu/pph2/), Sorting Intolerant from Tolerant (SIFT, http://sift.jcvi.org/), Protein Variation Effect Analyzer (PROVEAN, http://provean.jcvi.org), Mutation Taster (http://www.mutationtaster.org/), VarSome (https://varsome.com/), MutationAssessor (http://mutationassessor.org), and Combined Annotation Dependent Depletion (CADD, https://cadd.gs.washington.edu/). Variant interpretation was performed according to the 2015 American College of Medical Genetics and Genomics guidelines.

For the Polish case, whole-exome sequencing was conducted at the Institute of Human Genetics, Technical University of Munich, Germany, following established protocol [71]. Exome capture was carried out using the Twist Bioscience Exome 2.0 kit (Twist Biosciences, South San Francisco, CA, USA), and sequencing was performed on an Illumina NovaSeq 6000 platform to generate 150-bp paired-end reads. Sequence reads were aligned to the human reference genome GRCh37/hg19 using the Burrows-Wheeler Aligner (BWA). Single-nucleotide variants and small insertions/deletions were called using GATK HaplotypeCaller according to best-practice workflows. Bioinformatic processing and variant annotation were performed using EVAdb (Exome/Genome Variant Annotation Database; Munich, Germany). Rare variants were prioritized using a minor allele frequency threshold of <0.001 in gnomAD v4.1.0 and in-house sequencing databases, with adequate sequencing support defined as coverage ≥20× and variant allele fraction ≥20%. Candidate variants were visually inspected using the Integrative Genomics Viewer (IGV; Broad Institute, Cambridge, MA, USA). *De novo* status was confirmed by the absence of the variant in both parental datasets and verified by Sanger sequencing. Variant annotation and classification followed established procedures, including assessment of population frequency, evolutionary conservation, and *in silico* pathogenicity predictions, and cross-referencing against relevant disease and variant databases.

### Human fetal brain tissue acquisition and immunohistochemistry

The human fetal brain tissue was collected from voluntary surgical termination of pregnancy at CHR Hospital-Site Laveu, Liege, Belgium. Samples were obtained with informed, written patient consent in compliance with General Data Protection Regulation (GDPR) and the ethical principles set out in the “Declaration of Helsinki” and the “Good Clinical Practice”. The ethical protocol was approved by the Liège Hospital-Faculty Ethics Committee (ethical agreement number: 2021/216). Embryo gestational age was estimated by the transabdominal ultrasound and confirmed by measuring tibial length. The human tissue was collected directly after surgery and transferred to the laboratory within 1 hour in 95% O2/ 5% CO2 oxygenated artificial cerebrospinal fluid (aCSF, 85 mM NaCl, 2.5 mM KCl, 7 mM MgCl2, 0.5 mM CaCl2, 1 mM NaH2PO4, 26 mM NaHCO3, 10 mM D+-glucose, 50 mM sucrose). Brain tissue was fixed in 4% paraformaldehyde (PFA) at 4 °C overnight, cryoprotected in 30% sucrose, embedded in O.C.T. compound, and sectioned into 20 μm-thick sections using a cryostat (Thermo Fisher Scientific). Sections were stored at −80 °C until use. For immunostaining, sections were washed three times for 10 min each in 0.05% PBST and subjected to antigen retrieval using Dako Target Retrieval Solution (10× diluted) at 90 °C for 10 min. Sections were then washed three times for 10 min each in 0.05% PBST, followed by blocking in blocking solution (PBS containing 0.1% Triton X-100, 0.2% Tween-20, and 10% normal donkey serum, NDS) for 1 h. Sections were incubated with primary antibodies diluted 1:10 in blocking solution at 4 °C overnight. The following primary antibodies were used: rabbit anti-KIF2C (1:1000; Proteintech Group), mouse anti-SOX2 (1:300; BD Pharmigen), and rat anti-CTIP2 (1:300; Abcam). After three washes in 0.05% PBST, sections were incubated with secondary antibodies diluted 1:10 in blocking solution at room temperature for 2 h. The secondary antibodies used were as follows: donkey anti-mouse, anti-rabbit, or anti-rat Alexa Fluor-555, Alexa Fluor-488, and Alexa Fluor-647 conjugated antibodies (1:500; Invitrogen). Nuclear staining was performed using 4’,6-diamidino-2-phenylindole (DAPI) at room temperature for 15 min. All steps were carried out in the dark. Coverslips were mounted using antifade mounting medium. All images were acquired using a confocal laser microscope Zeiss LSM980 (Carl Zeiss).

### Plasmid construction for human *KIF2C*

Human *KIF2C* cDNA was synthesized based on NCBI reference sequence NM_006845 by Eurofins. The open reading frame (ORF) of human *KIF2C* was cloned into the *pCAX* vector using the In-Fusion cloning system (TaKaRa). To generate a vector to express disease-associated *KIF2C*, mutagenesis was performed by a modified QuikChange II site-directed mutagenesis protocol (Agilent Technologies) using KOD Plus DNA polymerase (Toyobo), followed by DpnI (NEB) digestion. Primers were designed at QuikChange Primer Design site (Agilent Technologies). The point mutation was confirmed by Sanger sequencing.

### Image analysis and quantification

Cell counts and fluorescence intensity measurements were performed using Fiji/ImageJ software (National Institutes of Health). For γ-H2AX intensity analysis, γ-H2AX intensity in GFP+ cells was measured on a per-cell basis and normalized to the DAPI intensity of the same cell. Fabp7 intensity was quantified as described previously [70]. Briefly, the mean fluorescence intensity within the region of interest was calculated after subtraction of the mean background intensity.

The length of the longest neurite in N2a cells was measured using Fiji/ImageJ. A line was drawn from the DAPI-labeled nucleus to the tip of the longest neurite, and the length of this line was defined as the neurite length. For Sholl analysis, image stacks of N2a cells immunostained with an α-tubulin antibody were traced and reconstructed as 2D binarized images. The center of each cell body was selected, and the number of intersections with concentric circles of increasing radii was quantified using the ImageJ Sholl analysis plugin [72].

### Statistical analyses

Statistical analyses were performed using GraphPad Prism v9.5.0. Two-tailed Student’s *t*-tests, multiple unpaired *t*-tests, and two-way ANOVA followed by Bonferroni’s or Tukey’s multiple-comparisons tests were used as appropriate. Differences were considered statistically significant at *p* < 0.05. Statistical significance is indicated as follows: \**p*<0.05, \*\**p*<0.01, \*\*\**p*<0.001, \*\*\*\**p*<0.0001.

---

**Supplementary Figure 1.**
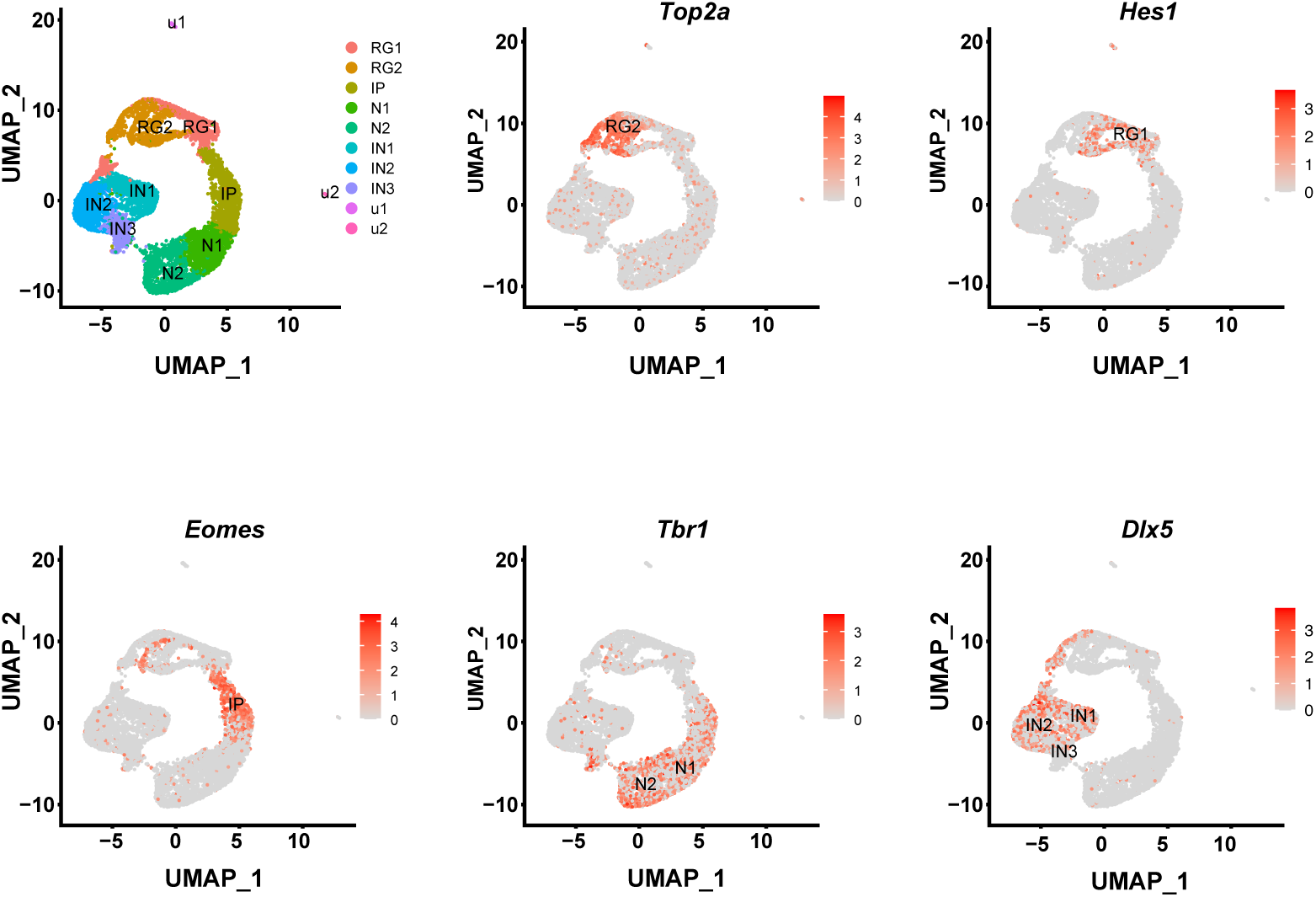
UMAP and feature plots of canonical markers for RG2 and associated cell populations, related to Figure 1. UMAP visualization of the E14.5 mouse cortical single-cell RNA-seq dataset identifies 11 distinct cell clusters, including two radial glia clusters (RG1 and RG2). IP, intermediate progenitor cell cluster; N1 and N2, neuron clusters 1 and 2; and IN1, IN2, and IN3, interneuron clusters 1, 2, and 3. Feature plots show the expression of canonical markers used to define cluster identities. The progenitor clusters RG1 and RG2 are characterized by expression of *Hes1,* whereas the proliferation marker *Top2a*, is specifically enriched in RG2. *Eomes, Tbr1,* and *Dlx5* mark intermediate progenitors, excitatory neurons, and interneurons, respectively.

**Supplementary Figure 2.**
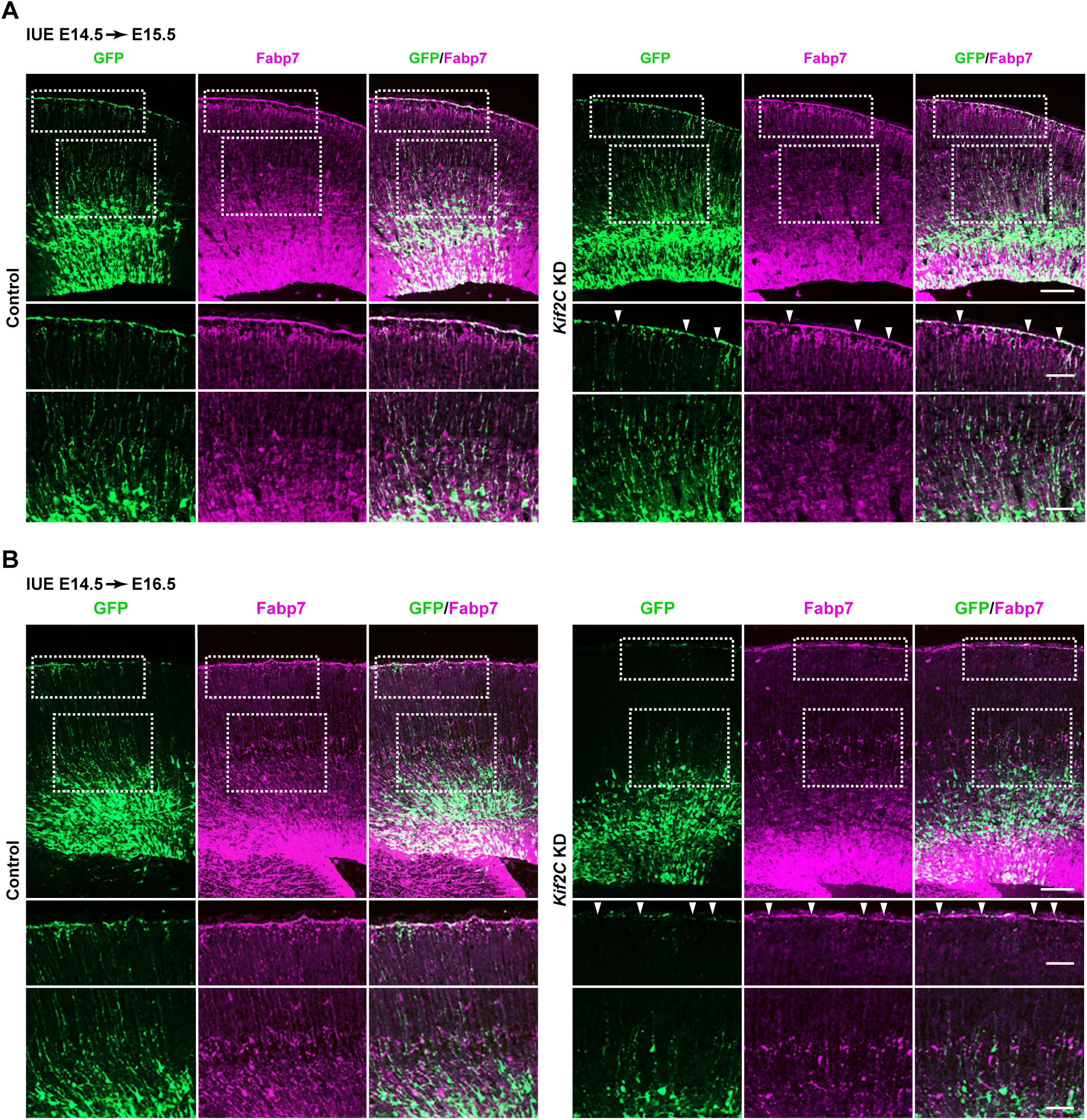
*Kif2C* knockdown causes progressive disruption of the radial glial scaffold, related to Figure 2. **(A, B)** Representative images of E15.5 **(A)** and E16.5 **(B)** mouse cortices, corresponding to 1 day and 2 days after in utero electroporation with control siRNA or *Kif2C* siRNA, stained for GFP and Fabp7. White arrowheads indicate GFP⁺/Fabp7⁺ basal endfeet. Boxed regions are shown at higher magnification in the lower panels. Scale bars, 100 μm (overview images), 50 μm (magnified images).

**Supplementary Figure 3.**
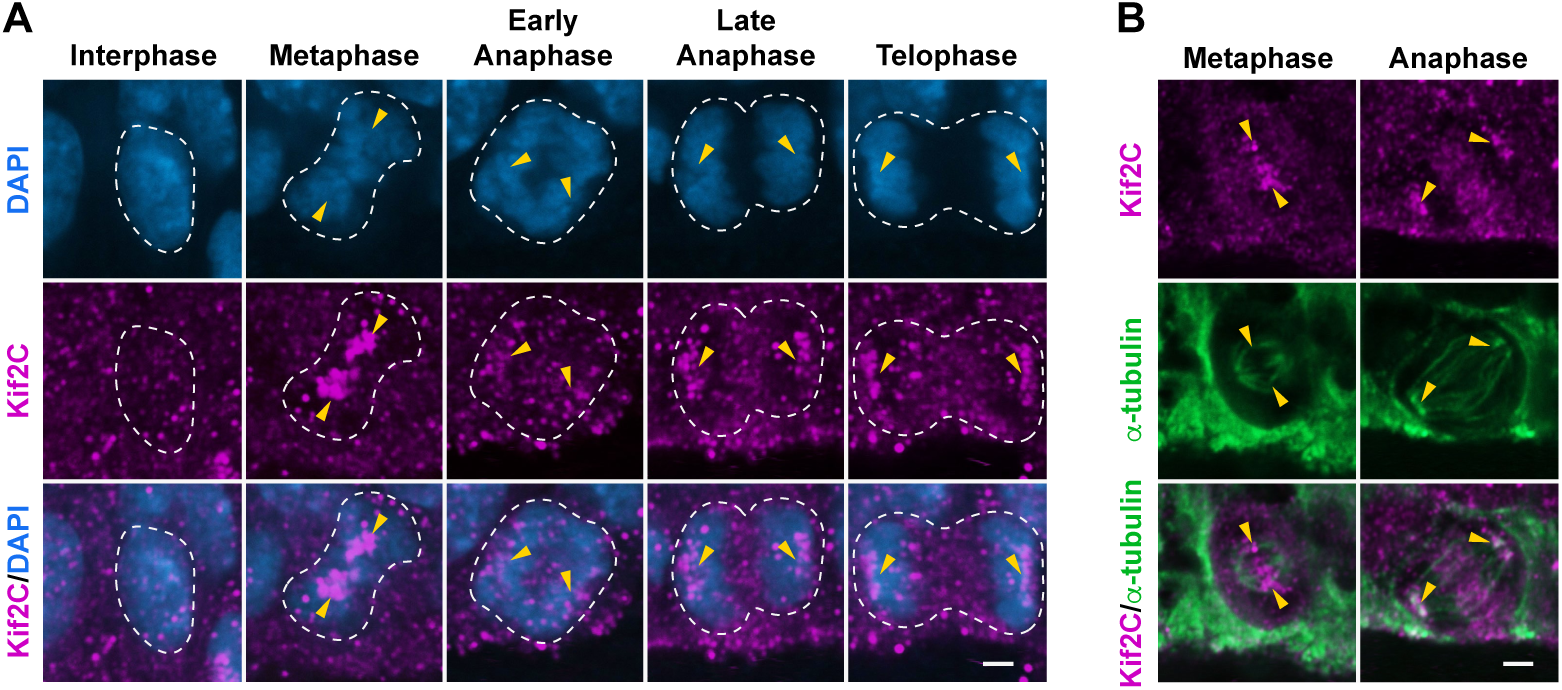
Subcellular localization of Kif2C in RGCs across the cell-cycle phases, related to Figure 3. **(A)** Representative images of WT E14.5 cortex stained for Kif2C and DAPI. Yellow arrowheads indicate Kif2C localization at different stages of the cell cycle. Scale bar, 2 μm. **(B)** Representative images of WT E14.5 cortex stained for Kif2C and α-tubulin (a microtubule marker). Yellow arrowheads indicate Kif2C localization at kinetochore-microtubule attachment site. Scale bar, 2 μm.

**Supplementary Figure 4.**
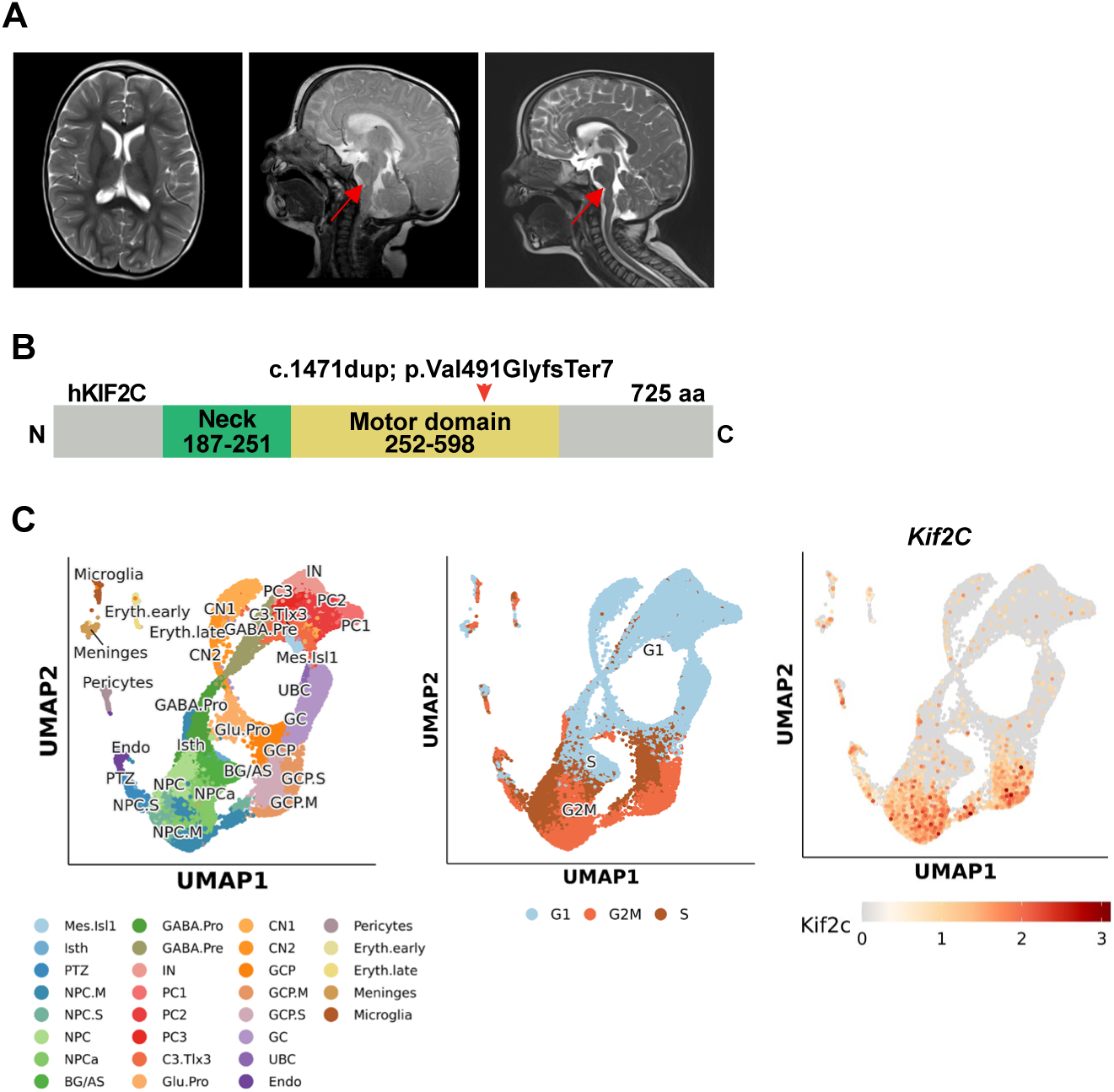
Phenotypic features of the Polish individual with a *de novo KIF2C* frameshift variant and enriched Kif2C expression in cerebellar progenitors, related to Figure 6. **(A)** Brain MRI images of the affected individual. Axial T2-weighted imaging (left) shows no visible supratentorial structural abnormalities. Sagittal T2-weighted images obtained at 5.5 months (middle) and 23 months (right) reveal abnormal brainstem morphology with hypoplasia, particularly at the pontomedullary junction (arrow). **(B)** Schematic diagram of KIF2C functional domains indicating the position of the identified *de novo* frameshift variant (c.1471dup; p.Val491GlyfsTer7). **(C)** Analysis of *Kif2C*expression in the developing cerebellum using a public single-cell RNA-seq dataset (Khouri-Farah et al., 2022). UMAP plots show cell-type clusters (left) and cell-cycle specific clusters (middle). Feature plots (right) demonstrate enriched *Kif2C* expression within cerebellar progenitor populations during the S/G2/M phases of the cell cycle.

**Supplementary Figure 5.**
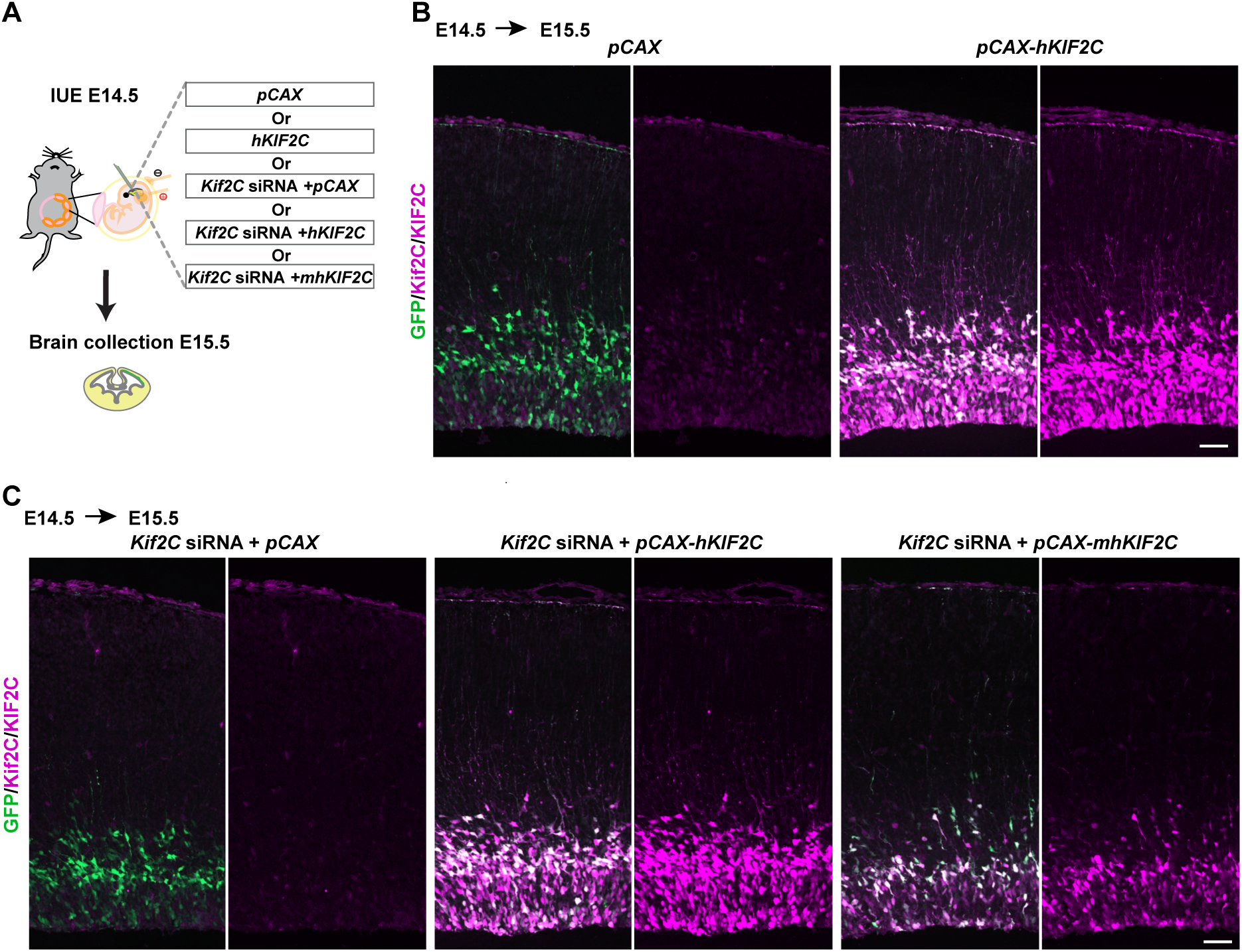
*Kif2C* siRNA depletes endogenous mouse Kif2C but not exogenous human KIF2C, related to Figure 6. **(A)** Schematic overview of the experimental design. **(B)** Representative images of E15.5 mouse cortices 1 day after electroporation with *pCAX* or *pCAX-hKIF2C*, immunostained for GFP and Kif2C/KIF2C. Scale bar, 50 μm. **(C)** Representative images of E15.5 mouse cortices 1 day after electroporation with control siRNA + *pCAX*, *Kif2C* siRNA + *pCAX-hKIF2C*, or *Kif2C* siRNA + *pCAX-mhKIF2C*, immunostained for GFP and Kif2C/KIF2C. Scale bar, 50 μm.

**Supplementary Figure 6.**
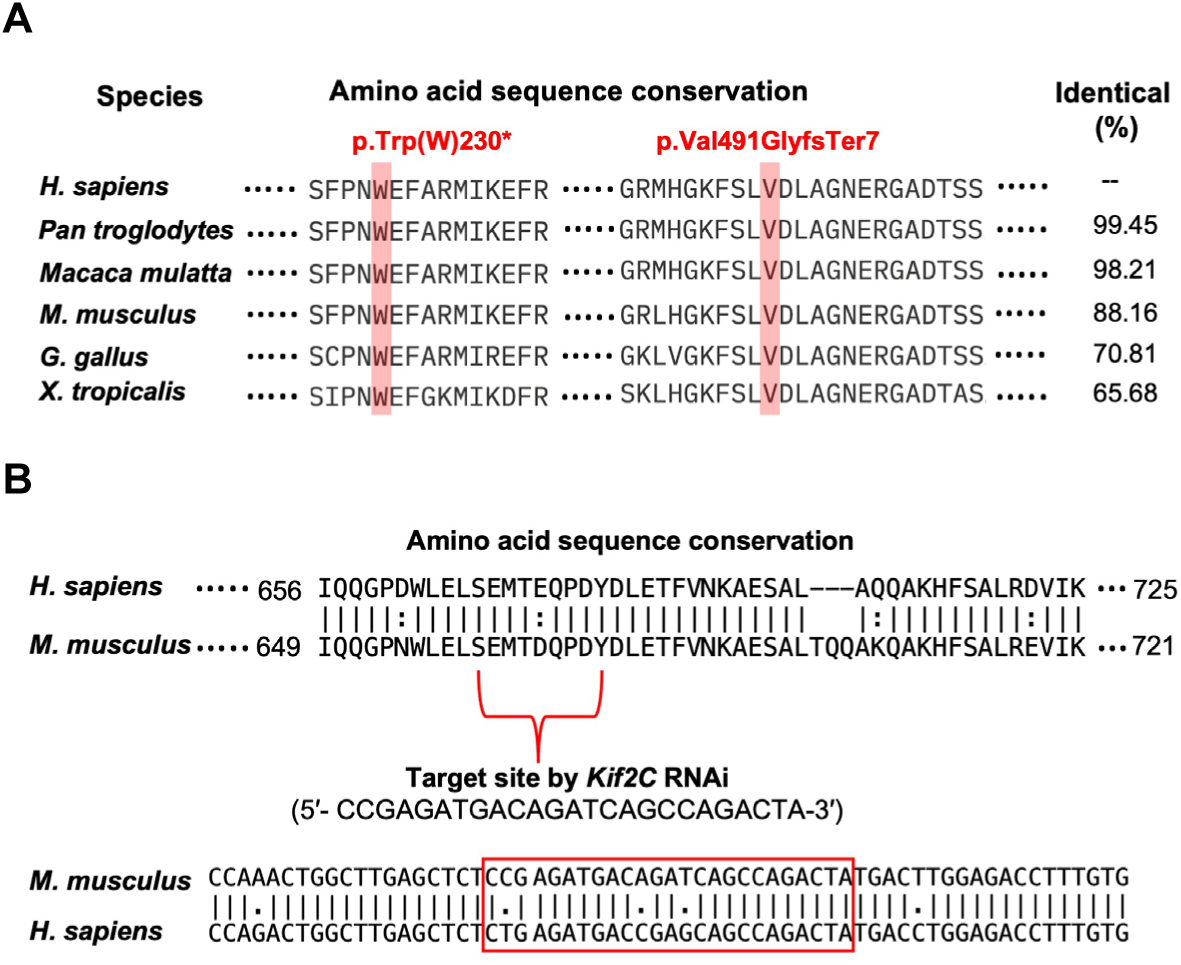
Conservation of Kif2C and mouse-specific RNAi design, related to Figure 6. **(A)** Multiple sequence alignment showing conservation of KIF2C across species. Red highlights indicate conservation of the regions affected by the clinical variants pTrp230* and p.Val491GlyfsTer7. **(B)** Alignment of human and mouse KIF2C/Kif2C amino acid sequences (top) and nucleotide sequences (bottom). The RNAi sequence was designed to match the mouse transcript while containing multiple mismatches with the human sequences (red box), thereby enabling mouse-specific knockdown.

**Table S1:** Frequency of cortical ectopia following *Kif2C* knockdown, related to Figure 5.

| **IUE E14.5 – E16.5** |  |  |  |
| --- | --- | --- | --- |
| Condition | Samples with ectopia | Total samples | % of samples with ectopia |
| Control | 0 | 14 | 0% |
| *Kif2C*-KD | 7 | 18 | ≈38% |
| **IUE E14.5 – P7** |  |  |  |
| Condition | Samples with ectopia | Total samples | % of samples with ectopia |
| Control | 0 | 4 | 0% |
| *Kif2C*-KD | 4 | 9 | ≈44% |

**Table S2:** Clinical features of patients with *KIF2C* mutation, related to Figure 6.

| **Catagory** | **Case 1** | **Case 2** |
| --- | --- | --- |
| Mutation | c.690G>A; p.Trp230* | c.1471dup; p.Val491GlyfsTer7 |
| Domain | Neck | Motor |
| Mutation type | Nonsense | Frameshift |
| Inheritance | Inherited | *De novo* |
| Sex/Age/Ethnicity | Male/22 years/Pakistani | Female/9 years/Polish |
| **Phenotypes** |  |  |
| Microcephaly | Significant (HC 49 cm, -4.11 SD) | Significant (HC 48 cm, -3.38 SD) |
| Neurodevelopmental status | Global developmental delay; intellectual disability | Global developmental delay; intellectual disability |
| Motor function | Delayed (independent walking at 19 months) | Severe delay (unable to sit independently) |
| Muscle tone/ Movement | Dystonia; impaired gross and fine motor coordination | Axial hypotonia; spastic tetraparesis, limb dystonia |
| Language/Speech | Impaired | Nonverbal |
| Neurological signs | Epilepsy (Onset 1.5-4 years; managed with Valproic acid) | Bilateral pyramidal syndrome (spasticity, hyperreflexia, extensor plantar reflexes, ankle contractures) |
| Sensory | Not specified | Congenital sensorineural deafness |
| Ocular/vision | left-sided ptosis; hypertelorism; downslanting palpebral fissures | Poor eye contact; inability to follow objects visually |
| Physical/ Dysmorphic | Reduced forehead; thick eyebrows, thin upper lip; midface hypoplasia, pointed chin | Thick eyebrows, epicanthus, short and narrow palpebral fissures, thin upper vermilion, pointed chin |
| Brain Imaging | Not available | Brainstem hypoplasia involving the lower pons, medulla oblongata, and pontomedullary junction |

